# Excessive immune responses in preterm placental villi compromise trophoblast health

**DOI:** 10.1101/2025.09.20.677440

**Authors:** Tyler A. Rice, Yi Cao, Leon Bichmann, Mar Ivar Ribes, Taylor A. Snisky, Serrena Singh, Weihong Gu, Dean Yimlamai, Jonathan S. D. Radda, Siyuan Wang, Raffaella Morotti, John S. Tsang, Christina J. Megli, Liza Konnikova

## Abstract

Appropriate regulation of maternal immune cells in the pregnant uterus is essential for successful pregnancy, yet the impact of immune cells in placental villi (PV) on placental health remain understudied. We enriched immune cells from term and preterm PV for transcriptomic and functional analyses. Maternal and fetal immune cells were abundant, with maternal chimerism increasing at term. Fetal lymphocytes displayed enhanced prostaglandin-D signatures, while maternal lymphocytes were enriched for antigen presentation pathways. Inflammatory cytokines and expanded maternal and fetal T cell clones peaked at term but did not induce cytotoxicity in placental organoids, whereas expanded T cells from idiopathic preterm cases impaired trophoblast growth and endocrine function *ex vivo*. These findings reveal significant immune chimerism within PV in both health and disease yet implicate PV T cells in idiopathic preterm birth.

**One Sentence Summary:** Fetal-maternal immune chimerism is characteristic of all placental villi, yet unchecked T cells and cytokines can hinder trophoblast growth and potentially precipitate preterm birth.

## Main Text

Successful human pregnancy depends on the enforcement of immune tolerance, such that neither maternal nor fetal immune cells reject the other, like rejection of a transplanted organ. Several mechanisms underlie this tolerance including maternal uterine NK cell (uNK) instruction of invading fetal extravillous trophoblasts, epigenetic silencing of class II MHC antigen presentation, and induction of suppressive regulatory T cells. (1) (2) (3) Less is known, however, about mechanisms regulating fetal immune cell reactivity. Fetal immune cells play an active role in development: yolk sac-derived fetal macrophages of the placenta, Hofbauer cells (HBC), are essential for supporting primitive hematopoiesis and placental function (2) (4) while mature, activated T cells in fetal intestines and lymph nodes respond to commensal bacterial antigens. (5) (6) (7) (8) Children born preterm suffer from wide range of sequelae include cerebral palsy, chronic lung disease, neurodevelopmental disorders, and increased risk of inflammatory diseases, yet two-thirds of preterm birth (PTB) cases are idiopathic and arise from unknown causes. (9) (10) (11) (12) Nonetheless, fetal immune cells can provide valuable insight into the effects of prenatal exposures during an early window of opportunity in which trajectories of future health and disease are being established. (13) Previous efforts to characterize fetal immune status in PTB have relied on umbilical cord blood, revealing phenotypes that are quite distinct from those in other tissues. (10) (14) Thus, we turned to the placenta as a source of tissue-resident immune cells that are central to tolerance, host-pathogen defense, other physiological functions, and form the basis of postnatal immunity extending into childhood and adulthood.

While the placenta and fetal membranes (the maternal-fetal interface, collectively) are comprised of several distinct tissue compartments, we chose to focus on placental villi, PV, and decidua basalis as the site of most direct exchange between mother and fetus. Several other groups have previously assembled invaluable single cell atlases of human placenta, but they have either been restricted to early gestation or have been quite sparse in adaptive immune cells, precluding in-depth examination of maternal and fetal lymphocytes and their functions. (15) (16) (17) (18) (19) (20) (21) (22) (23) (24) Dysregulated maternal T cells and granulocytes have been implicated in at least two complications of pregnancy: villitis of unknown etiology (VUE) and preterm premature rupture of membranes (PPROM), but we wondered whether immune dysregulation was a feature of PTB more broadly or whether adaptive immune responses are characteristic of normal gestation. (18) (25) To examine the role of placental immune cells during third trimester, we implemented a battery of single cell transcriptomic analyses and functional experiments including generation of a novel immune-trophoblast organoid co-culture system using placental samples from healthy term pregnancies and those from preterm births with diverse complications. Surprisingly, we found many fetal and maternal immune cells (hereafter, immune chimerism) in PV from both uncomplicated term and preterm cases. Beyond the lymphocyte-dense decidua, we report that PV also contain an active immune cell compartment, with diverse cell lineages, transcriptional states, and functional interactions with immune and non-immune cells that vary with gestational age and inflammatory milieu. These findings further our understanding of immune chimerism in the placenta and its dysregulation accompanying preterm birth.

### Generation of a placental immune cell atlas from term and preterm births

We assembled a case-control study of 32 placentas, all of which were dissected and viably cryopreserved within 12 hours of Caesarean delivery to avoid confounding effects of vaginal delivery (**Figure 1A**). We assigned placenta samples to one of four study cohorts to control for gestational age and diverse obstetric complications (**Table 1**). These included healthy term cases (mean gestational age, GA = 39.1 weeks), cases delivered preterm for maternal indications (structural complications) without inflammatory etiologies (mean GA = 35.5 weeks), cases of twins delivered preterm for fetal indications (transfusion syndromes) (mean GA = 32.7 weeks), and idiopathic preterm cases with diverse clinical courses (mean GA = 28.8 weeks). As expected, preterm infants from twin cases and idiopathic preterm cases had significantly lower birthweights than preterm structural cases and term cases. After thoroughly discarding maternal decidua tissue and collecting core PV just beneath the fetal membranes, tissues viably cryopreserved and then batched thawed where live immune cells were enriched from all samples and profiled by single cell transcriptomics. A subset of cases (N=28) was also evaluated using single nuclei sequencing in parallel from both PV and decidua basalis to capture cell types less amenable to live isolation (e.g. trophoblasts) (**Fig. 1A-1E**). Hereafter we will refer to nuclei sequenced and integrated with cell-based data as “cells” for the sake of brevity. Matched maternal blood (MB) and umbilical cord blood (CB) were also collected from two term dyads in order to provide sources of bona fide maternal and fetal immune cells, respectively. We sequenced each dyad separately, integrated all the data (>250,000 cells in total), and annotated cell clusters using a combination of canonical marker genes and de novo marker genes that emerged from this dataset using differential expression analysis (**Fig. S1A-S1B**). There were no statistically significant differences in the frequencies of major cell lineages in preterm placentas compared to term placentas (**Fig. 1E**) (**Fig. S2**). This demonstrates that our data were successfully integrated across different clinical sites, gestational ages, and tissue types (**Fig. S1C-S1D**), and that the preterm placental pathologies included in this study unlikely stem from gross deficits in immune development. In agreement with reports from other groups, trophoblast cells were captured at a much higher rate using single nuclei sequencing (**Fig. S1C**), which is an important complement to immune cell-focused approaches. (17) Cytotrophoblasts (CTB) and differentiated extravillous trophoblasts (EVT) and syncytiotrophoblasts (STB) were captured at equivalent rates across all study cohorts with no significant differences (**Fig S3A-S3B**). We also examined the impact of covariates like fetal sex, PPROM, and labor upon global transcriptional state and found that none drove significant batch effects (**Fig. S4)**.

**Figure 1.**
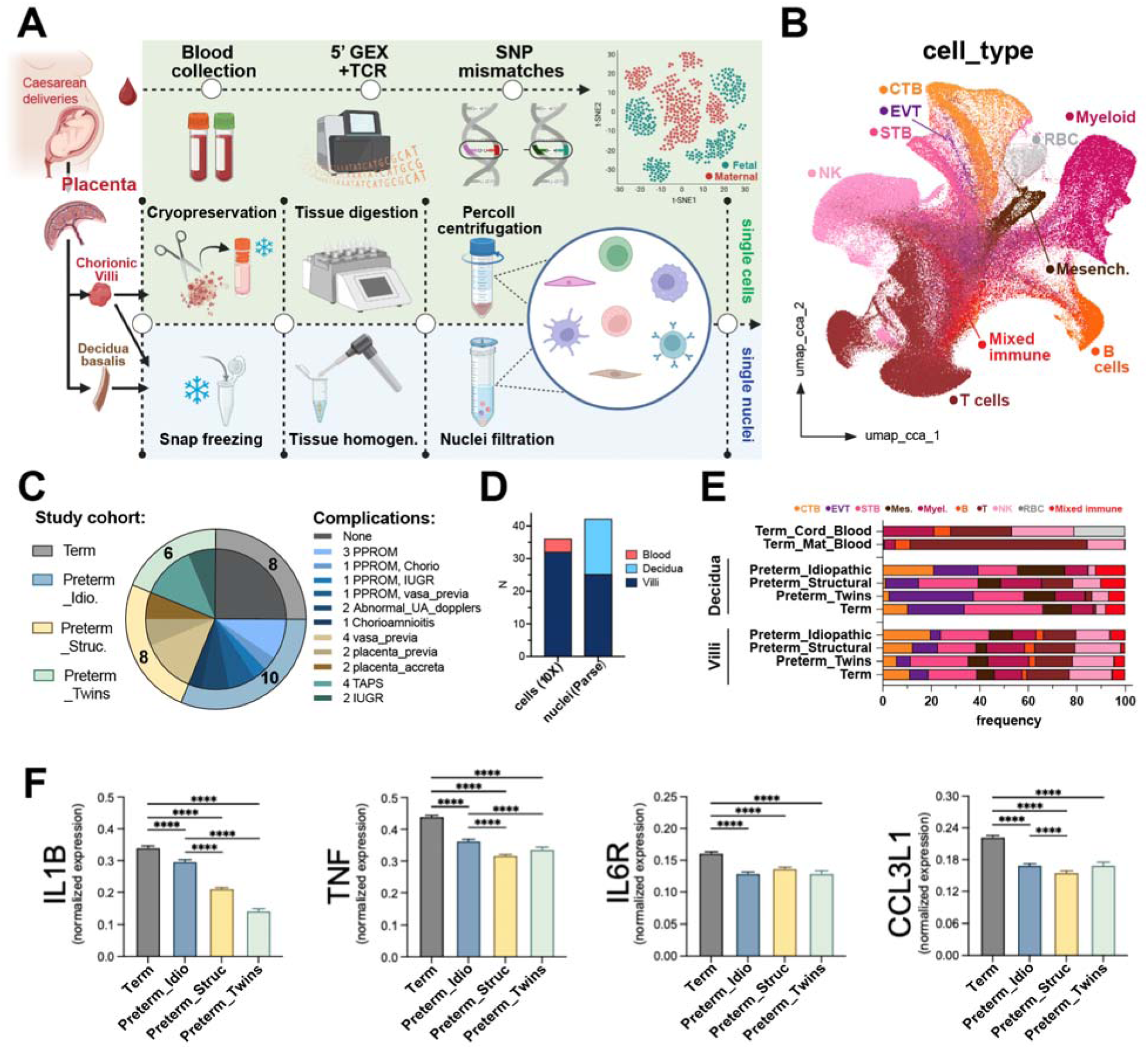
Generation of a placental villi (PV) immune cell atlas. (A) Schematic depicting sample collection, cryopreservation, tissue processing, isolation of cells and nuclei, and transcriptomic analysis methods. (B) Integrated placental immune cell atlas with annotated cell types encompassing ≥250,000 cells and nuclei. Abbrevs: CTB = cytotrophoblasts, EVT = extravillous trophoblasts, STB = syncytiotrophoblasts, NK = natural killer, RBC = red blood cells, Mesench. = mesenchymal cells. (C) Study cohorts and their constituent cases with complications of pregnancy. (D) Technology platforms utilized and sample type counts for each technology. (E) Frequencies of annotated cell types separated by tissue type (blood, decidua, and villi) and by study cohort (Preterm_Idiopathic, Preterm_Structural, Preterm_Twins, Term) (N=6-10 subjects/cohort). (F) Expression of cytokine-encoding transcripts across all placental villi cell types. \*\*\*\**P*<0.0001 by Kruskal-Wallis test.

### Both preterm and term placenta express inflammatory cytokines

We found strong inflammatory signatures across all cell types in term placentas prior to the onset of labor: high levels of cytokine-encoding *IL1B* and *TNF*, as well as *IL6R* and *CCL3L1* (**Fig. 1F**). Importantly, placentas from idiopathic preterm cases had higher *IL1B* expression than placentas from preterm twins or from structurally abnormal preterm cases, fitting with the clinical description of these cases (**Fig. 1F**). We found that trophoblasts from these idiopathic preterm cases expressed *CSH1* (encoding PL, placental lactogen) and long non-coding RNA *LINC02109* at levels trending higher than term cases (**Fig. S3C**), raising the question of whether this altered trophoblast transcriptional state resulted from normal development or PTB etiology. We also examined imputed cell-cell communication networks distinct across different cohorts and found that idiopathic preterm PV exhibited a mesenchymal cell signature between both immune cells and syncytiotrophoblasts (STB) (**Fig. S5**). Furthermore, while term PV adaptive immune cells both sent and received signals, preterm PV adaptive immune cells were predominantly receiving signals, suggestive of a dysregulated cellular network. Natural killer (NK) cells were highly implicated in many molecular signals, consistent with their known role in vascular remodeling during pregnancy and maternal tolerance. (15) Next, we sought to determine whether these immune cells were maternal or fetal in origin, using both tissue-level comparisons and cell-intrinsic computational approaches.

### A genome-wide demultiplexing approach reveals prevalent maternal and fetal immune cells, immune chimerism, in placental villi

First we used gold-standard approaches to examine the origin of villi immune cells: detection of both paternal-specific and maternal-specific HLA alleles (which were mutually exclusive in CB and MB) in PV immune cells (**Fig. 2A**), and visualization of XX+ HLA-DR+ maternal myeloid cells deep within core PV, far from the syncytiotrophoblast boundary, using in situ hybridization plus immunofluorescence (**Fig. 2B**) (**Fig. S6**). However, we sought a higher resolution approach that would reveal the origin of every individual cell within our PV immune cell atlas. Embryonic macrophages called Hofbauer cells (HBC) have long been described as the only PV-resident fetal immune cell type, in contrast to decidua maternal uNK cells and maternal T cells, which occasionally infiltrate villi during pathology. (25) To interrogate all immune cell types in PV rigorously, we used unsupervised single nucleotide polymorphism (SNP)-based demultiplexing to determine the fetal or maternal origin of each cell (**Fig. 2C**). (26) Trophoblasts served as an internal reference since they are exclusively fetal-derived by ontogeny. After overlaying cell type annotations, we found that 99.6% of trophoblasts were assigned to the same group. As such, all immune cells in the trophoblast-containing group were assigned fetal origin while cells in the other group were deemed to be of maternal origin (**Fig. 2D**). Each group was then benchmarked against blood: thousands of SNPs from fetal PV cells aligned strongly with SNPs from matched cord blood, while SNPs from maternal villi cells aligned with SNPs from matched maternal blood (**Fig S7A-S7C**). Beyond trophoblasts (**Fig. S7D**), demultiplexed fetal PV immune cells from male cases expressed Ychr-linked genes while demultiplexed maternal PV immune cells from the same male cases expressed Xchr-linked genes (**Fig. S7E**). Of note, the split-pool barcoding strategy used to sequence nuclei precluded determination of cell origins from any decidua or PV nuclei. We detected appreciable fetal myeloid cells, B cells, T cells, and NK cells and maternal myeloid cells, B cells, T cells, and NK cells across all 32 villi samples, rather than cell types restricted to exclusively mother or fetus (**Fig. 2E**). We were surprised to see that ∼30% of the *FOLR2-* and *CD163-*expressing myeloid cells in our single cell atlas were maternal in origin, which is inconsistent with the fetal ontogeny of HBCs (canonically expressing *FOLR2, CD163, FCGR2A* and *FCGR2B*). Marker genes reported to distinguish placental-associated maternal macrophages (PAMM) like *HLA-DRB1* failed to serve this purpose in our atlas since they were indistinguishable across true fetal HBCs and maternal HBC-like macrophages (**Fig. S8**). (27)

**Figure 2.**
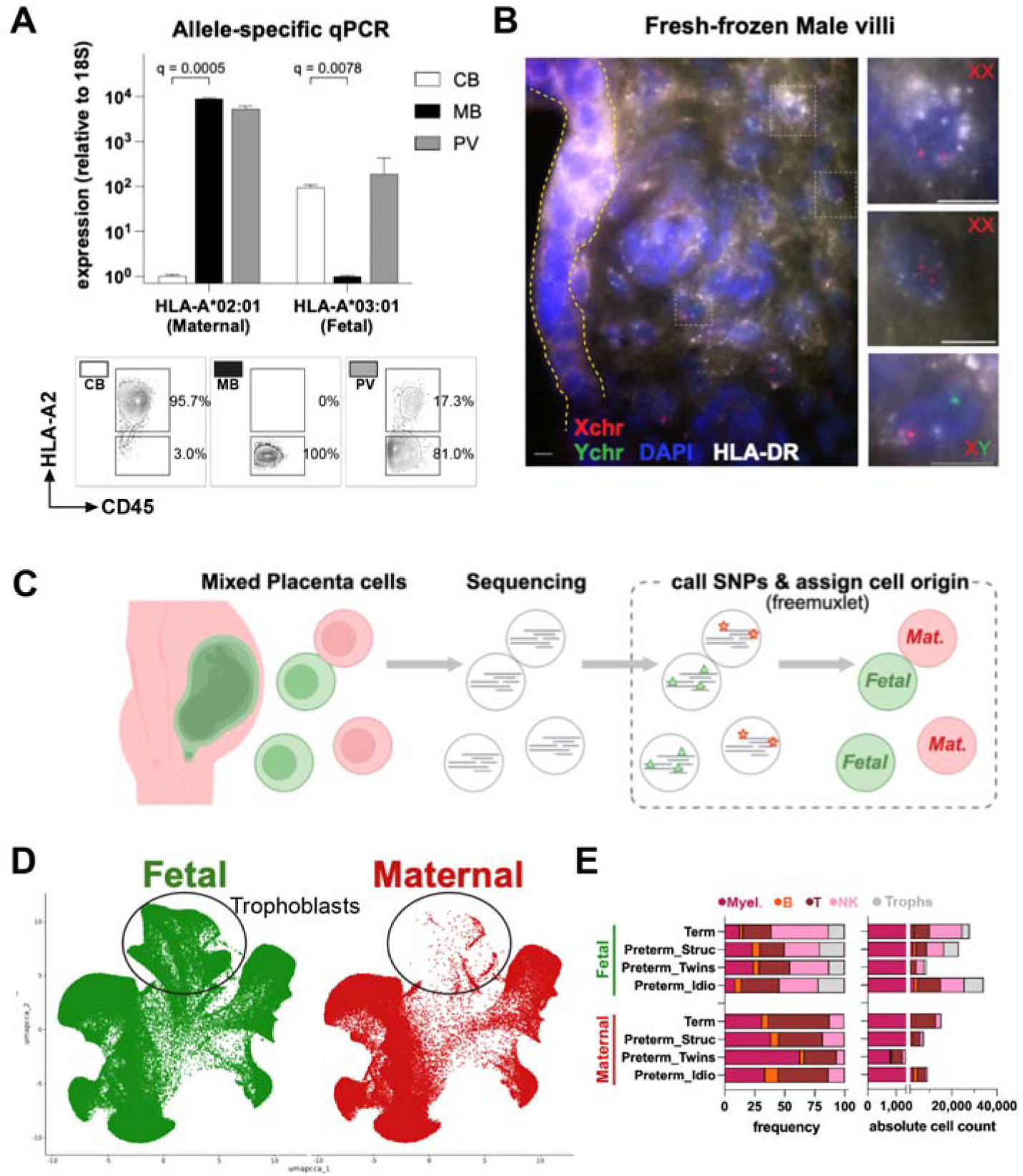
Both maternal cells and fetal cells are found in term and preterm placental villi. (A) HLA allele-specific qRT-PCR (top) and FACS (bottom) assessment of cord blood (CB), maternal blood (MB), and placental villi (PV) immune cells (N=1). q values generated by Mann-Whitney U test. (B) Micrograph (FISH + immunofluorescence) of male placental villi tissue section, stained for Xchr (red), Ychr (green), HLA-DR (white), and DAPI (blue) (N=1). Syncytiotrophoblasts marked by yellow dashes. Scale bars, 10µm. (C) schematic of SNP-based demultiplexing approach enabling discrimination of fetal or maternal origin. (D) Benchmarking cell origin discrimination on trophoblasts. (E) Frequencies and absolute counts of fetal origin and maternal origin cells across four study cohorts (N=6-10 subjects/cohort).

### Immune cell states in PV are distinct from both cord blood and decidua

Placental tissues are often discussed as cleanly segregated compartments: maternal immune cells in decidua and intervillous blood, fetal HBCs in PV, and fetal immune cells in cord blood. We carefully separated and cryopreserved each of these biosample types at two different institutions, then benchmarked them against one another using both traditional (tissue-specific) and innovative (origin-specific) approaches. Using pseudobulk analysis, we found that villi immune cells were distinct from both CB immune cells and from decidual immune cells (**Fig. 3A-3B**). While *CD44* and *STAT1* dominated the decidua immune signature, B cells were strongly enriched in PV-specific DEGs (*PAX5, IGHD, MS4A1, FCRL1*) (**Fig. 3A**). (8) Compared to cord blood T cells, PV T cells expressed more effector molecules *IFNG, CTLA4,* and *PDCD1* (**Fig. 3B**). These markers, plus *LAG3*, predominated in maternal PV T cells in contrast to fetal-specific *IGF2BP3* (**Fig 3C**). We next asked whether any of these signatures were specific to one of the etiologies of preterm birth found in our cohorts.

**Figure 3.**
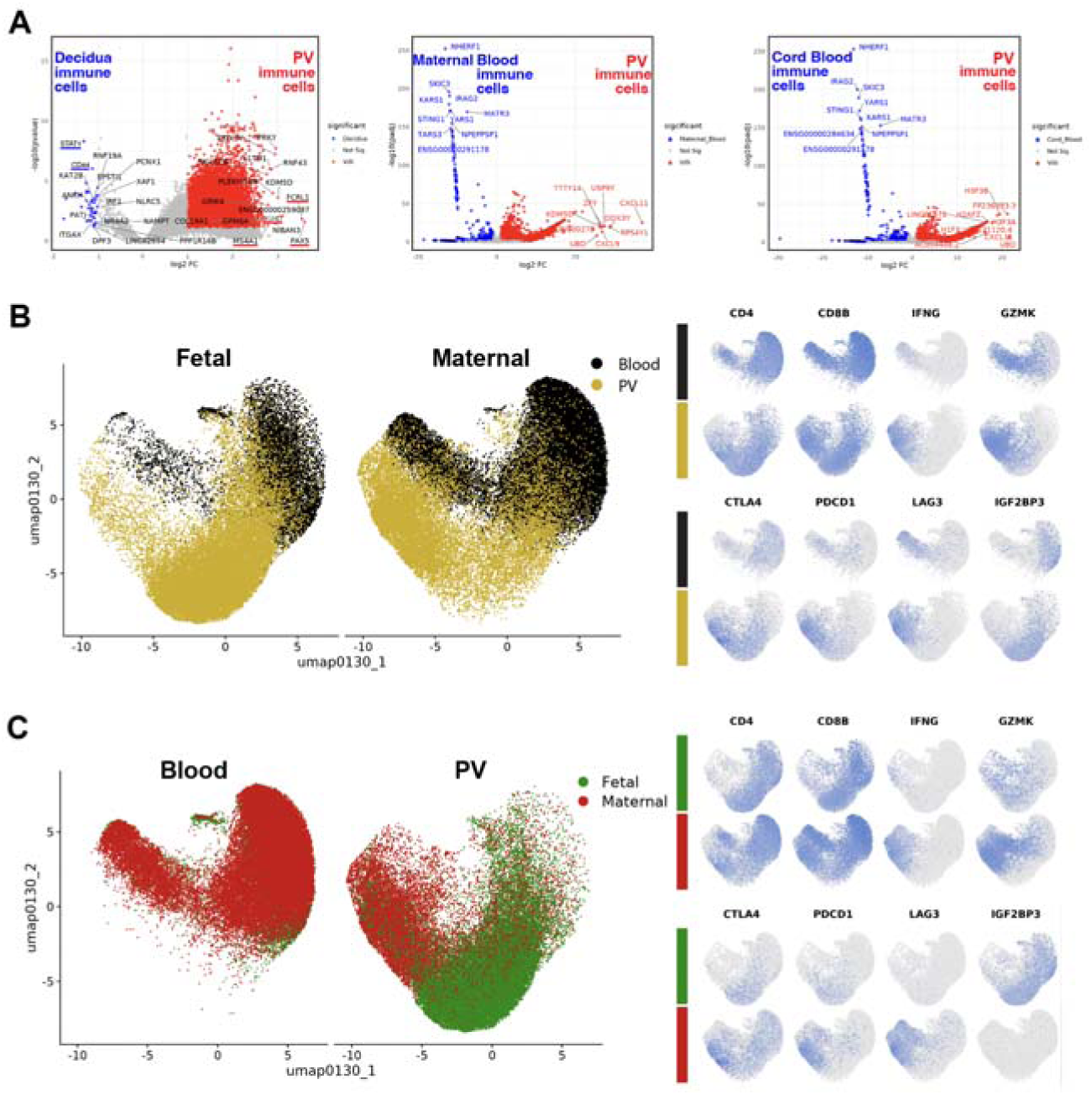
Tissue-specific and origin-specific features of PV immune cells. (A) Differentially expressed genes from pseudobulk analysis decidua, villi, cord blood, and maternal blood immune cells. (B-C) UMAP clustering and feature plots of T cells only demonstrating enrichment by tissue (B) and by origin (C).

### Preterm birth-specific and origin-specific gene signatures suggest partitioned functions

Prostaglandin D synthase (*HPGDS*) was enriched in myeloid cells while deoxyprostaglandin dehydrogenase (*HPGD*) was enriched in T and NK cells from idiopathic pretermcases (**Fig. S9A**). These data point to a potential dysregulation of fetal prostaglandin signaling in PTB. We also found that *FGL1,* a known ligand of co-inhibitory *LAG3*, was expressed more highly on decidual and PV macrophages from idiopathic preterm cases than term cases (**Fig. S9B**). (28) Meanwhile, antigen processing and MHC presentation gene signatures predominated in maternal immune cells, which were driven by term samples. (**Fig. S9C**).

### PV fetal immune chimerism decreases at term

We observed a consistent trend of decreasing frequency of PV fetal T cells, B cells, and Myeloid cells (not fetal NK cells) across gestational age, but not across fetal sex (**Fig. 4A-4C**). This led us to search for origin-specific features of fetal and maternal immune cells that might reveal unique developmental functions. We conducted differential expression analysis across maternal and fetal origin and found some origin-specific genes that may play roles in embryonic growth and programmed apoptosis (fetal *IGF2BP3*, maternal *AIM2*) (**Fig. S10A-S10B**). Despite the different environmental challenges faced by mother and fetus, cells of each origin seemed to converge on a highly similar transcriptomic state in PV, which perhaps hints at common goals. We also identified fetal-enriched *ROR1*, an orphan receptor with a known role in embryonic limb patterning using the Wnt pathway (**Fig. S10C**). (29) (30) At the protein level, we detected ROR1+ CD19+ fetal B cells by FACS in cord blood and PV, but not in maternal blood (**Fig. S10D-S10E**). The frequency of ROR1+ fetal B cells in PV samples was inversely proportional to gestational age, consistent with our assessments of immune chimerism based on single cell transcriptomics (**Fig. 4D**). We conclude that immune chimerism is a hallmark of PV across gestation with enhanced maternal contribution with gestational age progression, with diverse fetal and maternal immune cell types participating in normal embryonic development. To understand the contributions of specific immune lineages on placental health more deeply, we started with adaptive immune cells which have been far less studied in PV than macrophages.

**Figure 4.**
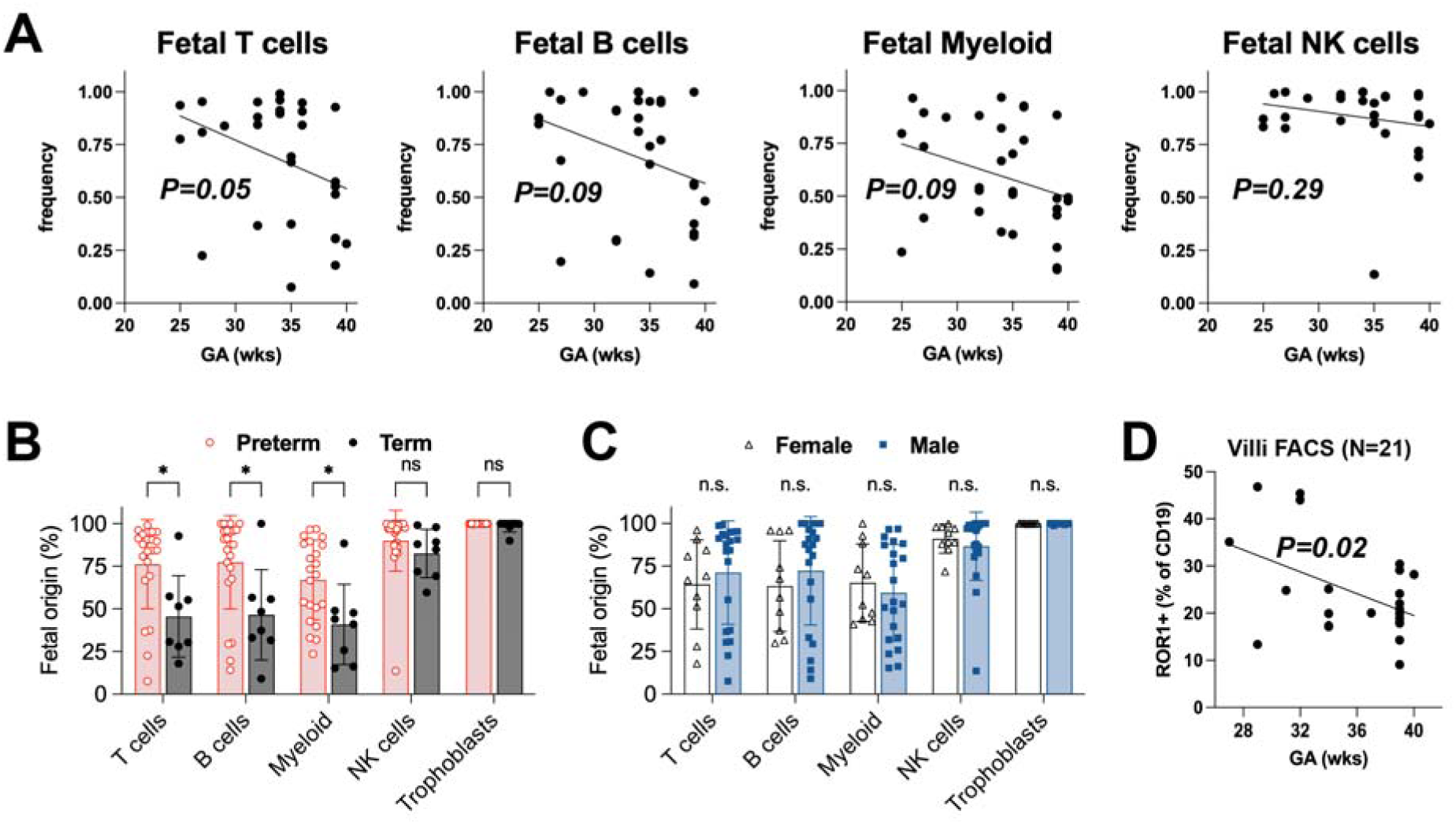
Fetal immune chimerism decreases in term PV. (A) Fetal immune cell frequency across gestational age (GA) (N=32). Fetal immune frequency by preterm status (B), or by fetal sex (C). \**P*<0.05 by Welch’s t-test. (D) Frequency of ROR1+ B cells in PV declined with gestational age (N=24). *P*-values from simple linear regression.

### Fetal B cells in PV are largely immature but display signs of early activation

Fetal hematopoiesis begins during first trimester, seeding mature B cells and T cells throughout many fetal tissues, including PV. (5) (6) (7) (9) (31) Compared to their maternal counterparts, PV fetal B cells exhibited less isotype class switching (**Fig. 5A**), which concords with their limited antigen encounters in utero. However, even though *CD38* was higher on fetal PV B cells, a known enforcer of early life fetomaternal tolerance, we still detected low levels of IgA+ and IgE+ fetal B cells and some that expressed memory marker *CD27* (**Fig. 5B**). (32) Compared to fetal B cells in cord blood, PV fetal B cells expressed more genes indicating early activation events (*NR4A1/2, CREM, FOSB,* and *DUSP2)* in contrast to the naïve blood signature (*TCL1A, LEF1*), once again demonstrating this uniquely active tissue compartment in contrast to cord blood (**Fig. 5C**). Next, we undertook a similar approach to examine PV T cells.

**Figure 5.**
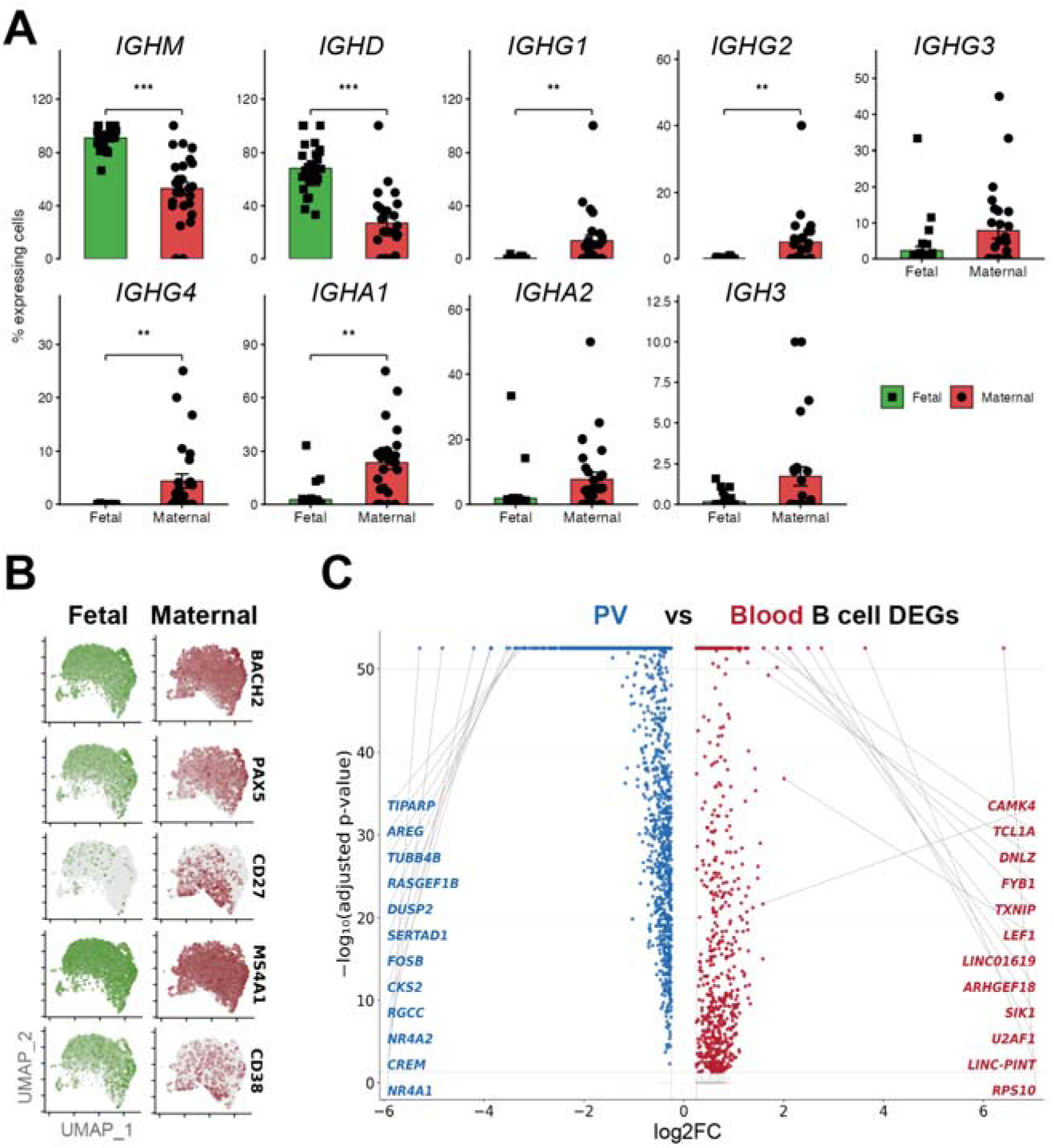
Maternal B cells in PV exhibit more isotype class switching and memory phenotypes than fetal B cells. (A) Per-subject isotype switching (N=32). \*\**P*<0.01, \*\*\**P*<0.001 by Wilcoxon rank-sum test. (B) Genes encoding core B-lineage transcription factors and immaturity (*CD38*) and memory (*CD27*) markers. (C) Differential villi signature versus blood B cell signature.

### PV-resident T cell receptor repertoires exhibit more clonal expansion with increasing gestational age

We captured the T cell receptor (TCR) sequences of more than 80,000 T cells from our PV atlas. TCR repertoires of term PV T cells were reduced in beta diversity compared to preterm TCR repertoires (**Fig. 6A**). These data align with our previous findings and those from other groups demonstrating that gestational age leads to selection of the fetal TCR repertoire and expansion of certain clones, resulting in reduced repertoire evenness. (7) (20) As expected, maternal blood TCR repertoires were greatly reduced in beta-diversity compared with cord blood (CB) TCR repertoires, consistent with pronounced T cell clonal expansion in adulthood (**Fig. 6B**). All PV repertoires, regardless of cohort, contained more expanded TCR clones than the CB repertoire, at a comparable level to maternal blood (**Fig. 6C**), although we found a high extent of “public” clonotypes shared across many subjects (**Fig. S11A**). The expanded subject-specific “private” clones were largely found among *IFNG*+ *GZMB*+ effector T (Teff) cells (**Fig. 6D**). Using GLIPH2, we found many variable TCRβ CDR3 amino acid motifs in the most highly expanded clones irrespective of gestational age, in contrast to prevalent polar serine and glycine residues in public clones (**Fig. S11B-S11C**). (33) To search for other more nuanced molecular signatures we utilized a large language model (LLM) trained on masked amino acids to evaluate our PV TCR repertoires. (34)

**Figure 6.**
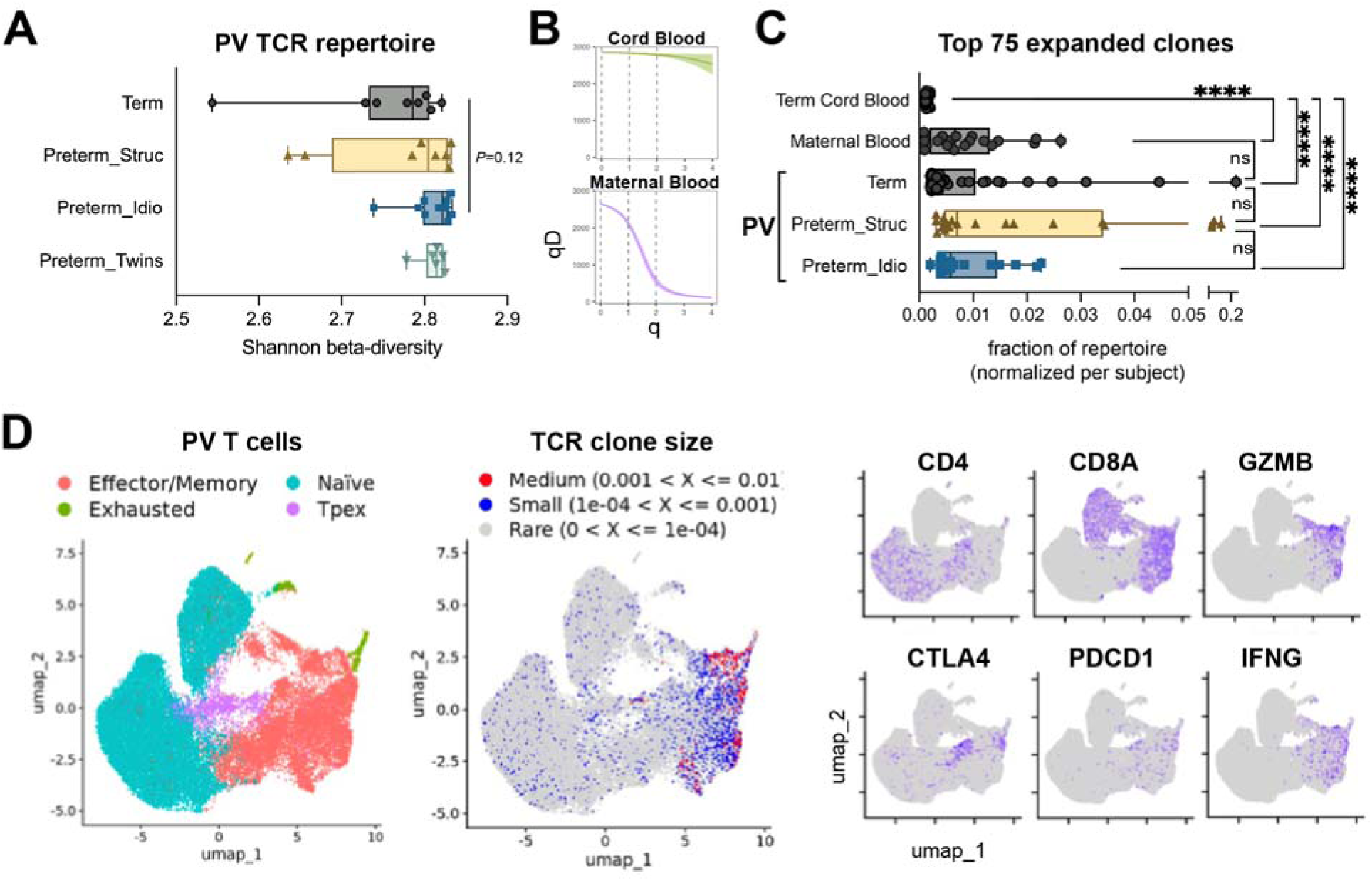
T cell receptor repertoires become more uneven with gestational age. (A) Shannon beta-diversity of PV TCR repertoires across study cohort (N=6-10). Box plots show interquartile range. P-val by Kruskal-Wallis test. (B) Hill diversity index, qD, of cord blood and maternal blood repertoires over increasing diversity orders, q. (C) Top 75 most expanded TCR clones split by study cohort. \*\*\*\**P*<0.0001 by K-W test. (D) Annotations of T cell phenotypes and TCR clonal expansion (left) and marker gene expression representing major T cell subsets (right). Abbrevs: Tpex = precursors to exhausted T cells.

### Preterm TCRβ motifs are robustly detectable using a large language model

Using TCRβ chain sequences from over 43,000 T cells, we classified clones that contained 3 or more unique T cells with an identical beta chain CDR3 sequence as “expanded”, while singletons and 2 unique cells we classified as “not expanded”. 70% of these data were used to train the TCR-BERT LLM, which was then tested on the remaining 30% (6,934 sequences) (**Fig. 7A**). Expanded clones were not robustly detectable by this model, so we performed the same classification exercise according to whether the TCRs originated from term or preterm PV. Leiden clustering of TCRβ sequences demonstrated that across a range of preterm and term predictions, each cluster of similar clones were scored uniformly rather than intermixed (**Fig. 7B**). TCR-BERT correctly detected preterm TCRs with much higher precision than term TCRs, with the sequences receiving the highest 5% preterm_score corresponding to a T cell population 81% of which derived from preterm PV samples (**Fig. 7C-7D**). This indicates that this model was both much better than random chance, and successful at detecting preterm TCRβ sequences across a wide range of criteria, not just a few expanded clones. Although LLMs often present challenges in extracting meaningful and straightforward factors that dictate the success of a given modeling task (interpretability), we found that the top scored preterm sequences contained biochemical motifs that were not detected by GLIPH2, including aromatic residues in positions 2, 4, and 11-14, and depletions of charged residues in positions 5-8 (**Fig. 7E**). These results suggest that prenatal inflammation may dysregulate the T cell compartment and aberrantly expand particular T cell clones. Next, we interrogated the functional effects of preterm PV T cells on trophoblast health.

**Figure 7.**
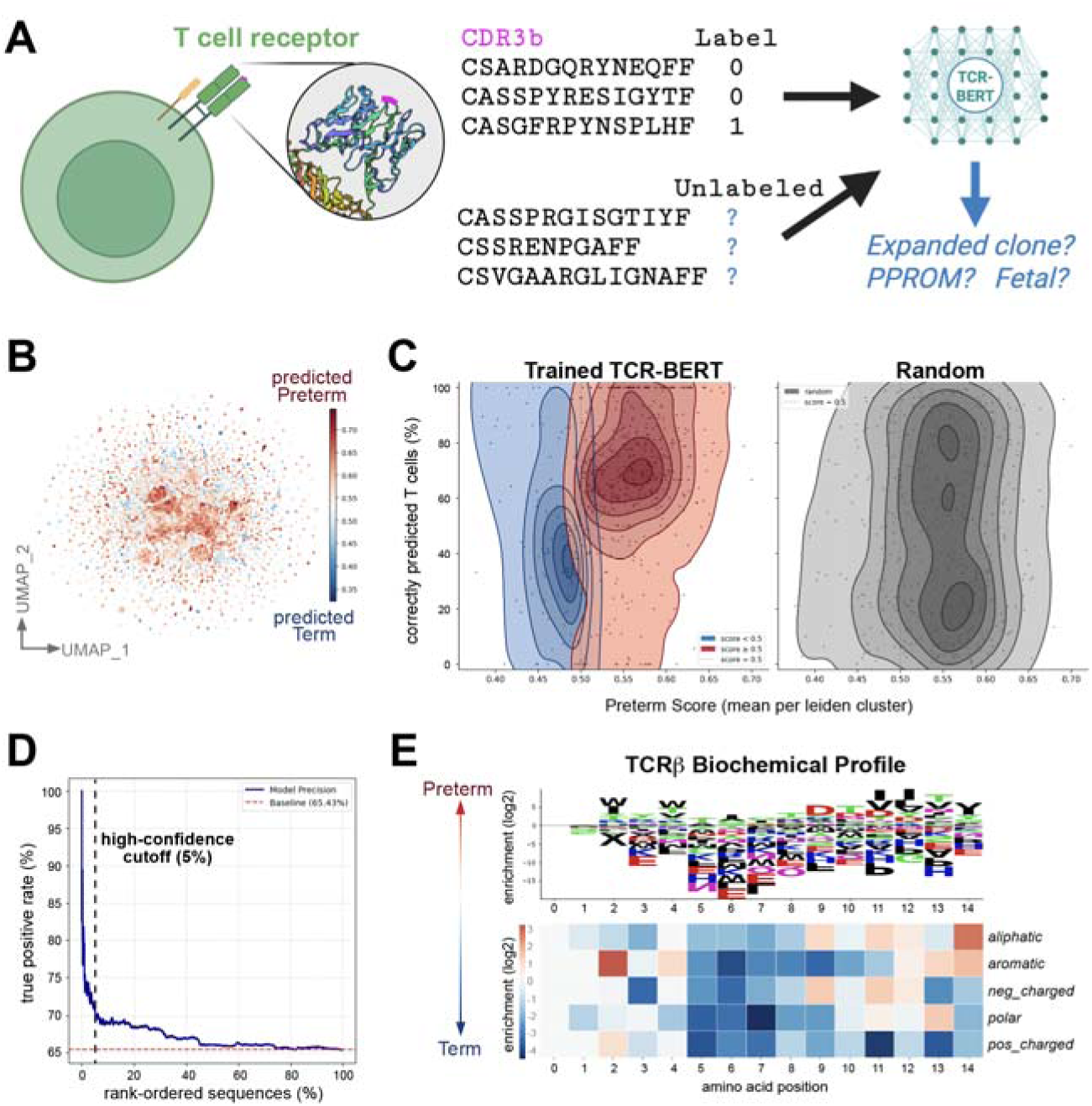
Large language model reproducibly detects preterm PV TCRs. (A) Training and testing approach using TCR-BERT. (B) Leiden clustering of TCRB embeddings (n=6,934 seqs), colored by preterm_score. (C) Prediction performance of TCR-BERT across preterm and term sequences, averaged by leiden cluster. (D) High-confidence detection of preterm TCRs in top 5% of preterm sequences. (E) Biochemical profiles of top 5% preterm TCRβ sequences compared to all term

### Preterm PV T cells but not term PV T cells impart inflammatory responses to primary trophoblast organoids in co-culture

To assess the functional contributions of PV T cells in term and preterm birth, first we cultured three-dimensional trophoblast organoids (TO) from term placental cases according to established protocols and found robust growth and hCGβ secretion (**Fig. 8A-8C**). (35) (36) (37) (38) (39) (40) Then we developed an immune cell–trophoblast co-culture system by co-encapsulating T cells from various sources within the same collagen gel as term TOs and measured organoid dynamics by light microscopy and molecular parameters. TOs co-cultured with T cells from idiopathic preterm PV exhibited slower growth and decreased KI67 and hCGβ (**Fig. 8C-8D**) (**Fig. S12**). This was not the case when we co-cultured T cells from term cord blood nor term PV, indicating that this is neither a global effect exerted by all T cells nor due to cytotoxic mediators perforin, granzyme B, or FasL, as these were below the limit of detection in both our RNAseq and Luminex assays. We performed cell type deconvolution on TO bulk RNAseq data and found approximately even representation of CTB, STB, & EVT trophoblasts across all co-culture conditions (**Fig. S13A-S13B**). However, the addition of idiopathic preterm PV T cells resulted in reduced expression of cell survival genes (**Fig. S13C**). We also compared TO bulk RNAseq with primary trophoblast single nuclei RNAseq, and found significant correlation of their transcriptional signatures, indicating that this is a representative model to maintain primary trophoblast gene programs in culture (**Fig. 8E**), which were not significantly skewed by fetal sex (**Fig. S14**). Co-culture of TOs with idiopathic preterm PV T cells led to elevated inflammatory cytokines IL-8 and MCP-1 in the supernatants compared to TOs cultured without T cells (**Fig. 8F**). These data lead us to conclude that although dysregulated preterm T cells do not kill primary trophoblasts outright (even in HLA-mismatched co-cultures), they can impair trophoblast growth and incite inflammatory responses *ex vivo*.

**Figure 8.**
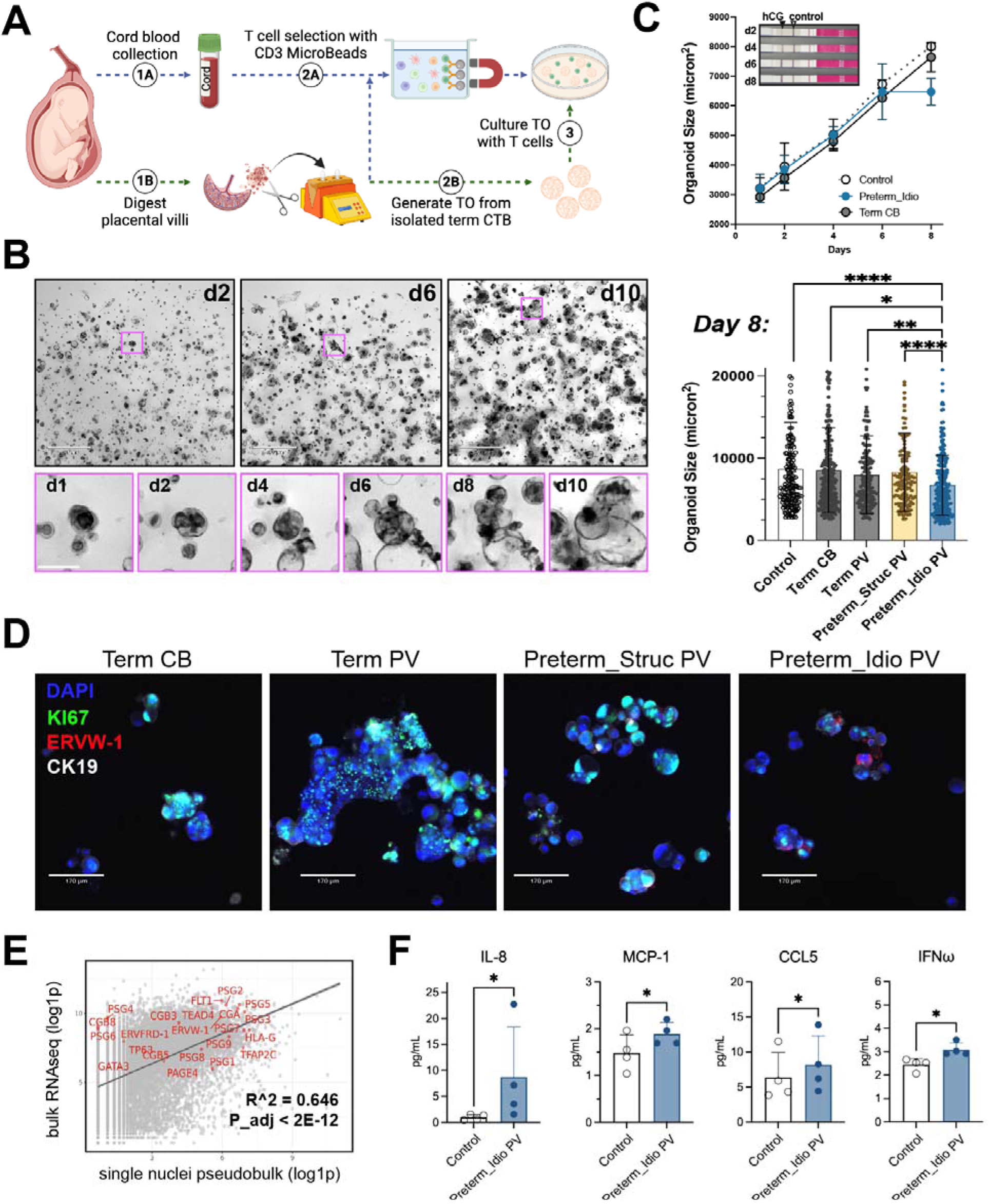
Preterm villi T cells dampen trophoblast organoid (TO) growth ex vivo. (A) Experimental schematic. (B) Brightfield images at 4X (top; scale bars 530µm) and insets of the same organoid growing over time (bottom; scale bars 100µm). (C) Organoid area measured longitudinally (top) and on d8 (bottom). \**P*<0.05, \*\**P*<0.01, \*\*\*\**P*<0.0001 by Krukal-Wallis test. (D) Immunofluorescence staining of d10 organoids. Scale bars, 50µm. Data shown is representative of N=3 independent experiments each with distinct TO lines. (E) Pearson correlation analysis across bulk and single nuclei RNAseq. (F) Cytokine levels in co-culture supernatants, assessed by Luminex. \**P*<0.05 by ratio paired t-test.

## Discussion

Many studies have detailed maternal immune cell tolerance in the decidua or fetal immune status in cord blood. (9) (10) (41) (42) (43) (44) However, less is known about the mechanisms of fetal immune development in the PV and cellular adaptations to allograft, infectious, or inflammatory insults. We sought to learn about fetal immune development in PV in greater detail in both health and disease. Using genome-wide SNPs to demultiplex maternal and fetal cell origin in PV, we have shown that not only decidua but also PV contain pervasive maternal immune cells, and that this is a distinct immune compartment. Immune chimerism extended to all cases examined, irrespective of pathology, PTB, or gestational age, although the magnitude of maternal chimerism increased in late gestation.

We hypothesize that maternal immune cells may migrate through anchoring PV, as the first trimester observations of PAMMs perhaps suggests, and exert pro-labor pro-detachment functions before labor onset. We contend these chimeric cells do not simply derive from contaminating intervillous blood during our tissue processing based on four lines of evidence: Firstly, PV-resident maternal T cells were transcriptionally distinct from maternal blood T cells, forming separate clusters in UMAP space (**Fig. 3B**). Secondly, routine histology of our PV sections shows intervillous spaces completely free of RBC, with intact STB boundaries. Thirdly, in male placentas that were fresh-frozen and FISH stained for sex chromosome, XX+ maternal cells were not located at the villous boundary, but rather deep within the villous core (**Fig. 2B**)(**Fig. S6**). Fourthly, in contrast to B cells, T cells, and Myeloid cells, the PV NK cell compartment was almost exclusively fetal across gestation, in the face of highly prevalent maternal uNK cells (**Fig. 4A**). These data lead us to hypothesize that there might be cell type-specific chemotactic cues that orchestrate precise positioning of fetal NK cells and maternal NK cells in distinct layers of the placenta (decidua, PV, and perhaps syncytial knots) to carry out niche-specific functions. We eagerly anticipate how spatial transcriptomics approaches will enable rigorous evaluation of this hypothesis. (24) (45)

In concordance with previous reports, we show that known clinical indicators of idiopathic preterm birth are indeed borne out at the molecular level with many different immune and non-immune cell types producing *IL1B* and *TNF*. We predict that placenta-specific transcription factor networks and epigenetic mechanisms are in place to regulate time-sensitive immune-mediated inflammation for the sake of successful pregnancy and parturition. Stated another way, we expect there to be high context dependence of these cytokine signals, with differential effects across pairs of “sender” and “receiver” cells.

In contrast to the view that strong adaptive immune responses in the placenta are only pathological, our data show profound T cell activation, clonal expansion, and presence of effector memory T cells in both healthy term pregnancies and in some cases of preterm births. What the precise functions of these T cell responses are, especially in normal development, remain unknown, but one possibility is that T cells are being tolerized to innocuous antigens (food, commensal microbes) coming from maternal circulation. We are eager to assess T cell antigen specificity using antigen affinity reagents and other approaches like TCR-BERT and immunopeptidomics that can predict or measure what peptide antigens might be presented to these fetal T cells. We hypothesize that early life immune education is critically important for dictating subsequent adaptive immune events in childhood and adulthood. Indeed, there is a deep literature about the tolerogenic nature of early life immune cells, and we are hopeful to examine how PV might support the functions of other tissues (thymus, GI tract) that similarly foster immune development and limit autoreactivity.

Three dimensional trophoblast organoids are a powerful tool to interrogate cellular and molecular interactions ex vivo, and the field has exploded in the last decade with many important advances. (35) (36) (37) (38) (39) (40) In this study we extend these systems further by co-culturing TOs with primary PV T cells from idiopathic preterm cases, which limited TO growth compared to TOs co-cultured with term villi or term CB T cells. Although preterm PV T cells did not induce TO cytotoxicity, the mechanism of dysregulation remains unclear. Co-culture of TOs with other immune cell types may enable further functional dissection, using approaches like blockade of cytokines and other ligand-receptor interactions. We also hope to explore the possibility of live cell imaging to capture dynamic cellular interactions in real time.

Some limitations of our study design include a lack of samples exhibiting common maternal comorbidities like pre-eclampsia. Many complications of pregnancy present as constellations of symptoms that likely stem from interwoven biological processes, making neat categorization of heterogeneous cases a great challenge. We were also unable to collect paired maternal blood and cord blood for all of our placenta samples, which would provide more rigorous paired identification of PV-specific gene signatures in each dyad. Ex vivo, our TO co-culture experiments lack physiological blood flow, gradients in oxygen tension, and nutrient/waste exchange, and are currently dependent on animal-derived extracellular matrix (Matrigel) in order to form 3D organoids. Lastly, we still lack the necessary technological advances to purify maternal PV T cells from fetal PV T cells that would allow us to parse their respective functions ex vivo.

Preterm birth has been a difficult problem to tackle for the research community, with only incremental improvements in efforts to decrease its prevalence in the last ten years. Identification of which etiologies of preterm birth are preventable (and how) remain unsolved, and we are interested in understanding which are immune-mediated. In this work we show that even some preterm cases delivered for maternal indications (preterm_structural) exhibited surprising levels of T cell clonal expansion (**Fig. 6A-6C**), demonstrating that the PV immune system is not necessarily quiescent in the absence of membrane rupture. Although these T cell responses are clearly maturing in both healthy term cases and preterm cases, were surprised to find that TCRβ sequences from expanded clones were not similar enough to one another to yield distinguishing features. Rather, the trained TCR-BERT large language model was able to detect features of preterm TCRβ sequences with high confidence. We wonder if this might indicate an immune-mediated origin of preterm birth that could be developed into a biomarker of PTB risk. Whether or not there is an upstream antigenic trigger or inherited genetic allele (e.g. HLA, etc.) leading to this T cell response remains to be examined in future studies.

Despite these challenges, preterm birth is an enormous risk factor for many comorbidities throughout childhood and adulthood, and we contend that this devastating condition necessitates further study. The restraint of alloreactive maternal T cells against the fetus is a remarkable immunological feat, and we intend to leverage TO-immune co-cultures to find the key factors mediating such immunoregulation. Immune-mediated preterm birth is a prime target for new strategies for patient risk stratification, and, in the era of widespread immunotherapy successes, translational lead development, enabling physician scientists to limit inflammation, delay labor, and augment childhood health in one fell swoop.

## Materials and Methods

### Study Design

Thirty-two placentas were evaluated in this study according to the following criteria

Inclusion criteria: pregnant patients admitted to labor & delivery unit, greater than 18 years of age, English-speaking. Exclusion criteria: smoking and recreational drug use, labor augmentation or induction, multiple pregnancies, fetal abnormalities, vaginal delivery, congenital anomalies. Medical history data was collected by obstetric providers, and no additional follow-up information or specimens were collected from these patients. There were no endpoints or replicates inherent to this study since it was neither interventional nor longitudinal. Our objective was to assess immune signatures in different modalities of preterm birth. To that end we assembled three different cohorts of preterm placentas (**Table 1**), all of which were compared to term controls (N=8): preterm_structural (N=8), delivered for maternal indications, preterm_twins (N=6), delivered for fetal indications (transfusion syndromes), and idiopathic preterm cases with diverse pathologies (preterm_idiopathic) (N=10). Researchers were not blinded to sample identity.

### Study approval

All protocols were approved by institutional review boards at Yale University (protocol #2000028847), or University of Pittsburgh (protocol #19100322), and all subjects were provided informed consent to participate in research.

### Placenta tissue procurement

Placental samples were obtained from three distinct biobanks. Placentas from cases with structural complications and twin transfusion syndromes were collected by the University of Pittsburgh Obstetrics & Gynecology. Placentas from preterm labor cases were collected by Yale New Haven Neonatology in coordination with Obstetric Pathology in cases being consulted for NICU admission. Term samples were collected from the Yale University Reproductive Sciences (YURS) Biobank after elective C-section deliveries scheduled prior to the onset of labor.

### Placenta sample collection, processing, & cryopreservation

After Caesarean delivery, each placenta was refrigerated and transferred to the laboratory in less than 12 hours. In a biosafety cabinet, placentas were dissected as follows, adhering to aseptic technique. Halfway between the umbilical cord insertion site and the edge of the basal plate, 2cm x 2cm full thickness cores were taken using sterile scissors, avoiding large fetal blood vessels. Maternal decidua was removed using a scalpel, then fetal membranes were removed, and chorionic villi were dissected from the basal plate, transferred to a Petri dish, and washed two times with sterile 1X PBS. Pieces of villi (∼0.5 cm^3) were placed either in histology cassettes and immersed in 10% neutral-buffered formalin for 24h or placed in cryomolds in room temperature OCT media and fresh frozen by floating atop liquid nitrogen. For generation of cell suspensions, villi were then incubated in 1X RBC lysis buffer (eBioscience #00433357) for 10min, then washed a further two times in 1X PBS before vacuum aspirating all traces of PBS and replacing with freezing media: heat-inactivated fetal bovine serum (FBS) +10% DMSO. Villi were minced with sterile scissors then transferred to 2mL cryovials (50-100mg tissue per vial) using a P1000 micropipet and wide-bore pipet tips. Cryovials were placed in a room temperature Nalgene Mr. Frosty containing isopropanol, which was then transferred to -80C freezer for at least 24 hours. Cryovials were then transferred to liquid nitrogen for long term storage.

### Cell isolation

Immune cells were enriched from placental villi (PV) as follows: Cryovials of minced tissue were thawed in 37C water bath for 2-3min, then tissue was transferred to 14mL 1X PBS, centrifuged 2min at 500xg to remove DMSO-containing media, which was aspirated and discarded. Tissue was transferred to 10mL digestion media (RPMI 1640 media, 10% FBS, 20mM HEPES, 1X L-Glutamine, 1X Pen/Strep, 1X Sodium Pyruvate, 1X Non-essential amino acids, and enzymes from Umbilical Cord Dissociation Kit (Miltenyi #130-105-737): 10µl enzyme A, 4µl enzyme B, 100µl enzyme D, and 62.5µl enzyme P) in an C-tube (Miltenyi #130-096-334), and placed into Miltenyi OctoMACS digestion instrument warmed to 37C for 1 hour of mechanical digestion. After digestion, an additional 10mL of complete RPMI was added, and cells were gently triturated using 10mL serological pipet and filtered serially through at least three filters: stainless steel mesh, 100µm nylon (repeating if necessary), and 70µm nylon. Cell suspension was centrifuged 650xg for 10min, resuspended in 10mL 25% Percoll (diluted in complete RPMI), and layered dropwise over 10mL 70% Percoll, down the side of an angled 50mL conical tube, being careful not to disturb the boundary between Percoll layers. Assembled percoll tubes were centrifuged 30min 800xg at 4C with acceleration set to 1 and brake off. Upper debris/trophoblast layer was discarded, and cells sedimented at the interphase were transferred to 10mL complete RPMI media, spun down at 650xg, resuspended in 1mL PBS/0.04%BSA, counted by trypan blue exclusion, and adjusted to a concentration of 1.6e6 cells/mL. If viability was <70%, cells were centrifuged and subjected to Dead Cell Removal kit (Miltenyi #130-090-101) using MS columns and collecting the live flow-through fraction.

### Library preparation & sequencing

Live cells were adjusted to a concentration of 1.6e6 cells/mL and then emulsified with RNA capture beads using the 10X Genomics Chromium controller. cDNA libraries were generated in tandem using 5P-GEM-X and 5P-VDJ reagent kits (10X Genomics). Quality of sequencing libraries was evaluated using Agilent Bioanalyzer, and multiplexed libraries were sequenced on Illumina NovaSeq S4 to a target read depth of 200M read pairs per sample.

### Nuclei isolation & sequencing

Snap-frozen placental villi and decidua tissues were homogenized right after removal from -80°C using a disposable pestle attached to a cordless motor in homogenization buffer (250mM sucrose, 25mM KCl, 5mM MgCl2, 10mM Tris buffer, pH 8, 1uM DTT, 0.16U/ul RNase inhibitor, 0.1% DTT) and DNase. Sample was then mixed with additional homogenization buffer and incubated on ice for 5 minutes. Homogenate was filtered through a 40mm strainer. 500K nuclei were transferred to a new DNA LoBind tube for fixation. Nuclei fixation was carried out using Evercode™ Nuclei Fixation (Parse Bioscience). Barcoding and library generation was performed using Evercode™ WT (Whole Transcriptome) v2 kit (Parse Bioscience). Pooled libraries were sequenced on Illumina NovaSeq S4 to a target read depth of 200M read pairs per sample.

### Bioinformatic analysis

Sequencing reads were aligned to the human transcriptome assembly GRCh38-2020a and assembled into count matrices using cellranger count. Matrices were then pre-processed as follows: batch correction (BBKNN), removal of doublets (doubletfinder), removal of ambient RNA (soupX), filtering on cells with >250 nFeatures captured and <25% mitochondrial gene content. Downstream analyses, including normalization, scaling, integration, clustering, annotation, and plot generation were all conducted in Seurat (v5.1.0). (46) Receptor-ligand analyses were conducted using CellChat. (47) T cell receptor repertoires were analyzed using nf-core/airflow. (48) (49) CDR3 sequences were then labeled according to their expansion size, preterm/term origin, or PPROM status, and used as input to the embed_and_classify function within TCR-BERT (34).

### Reference-free demultiplexing of maternal and fetal cells

The filtered barcoded single cell data was demultiplexed using the nextflow pipeline hadge. (50) Freemuxlet was selected as reference-free tool within this pipeline, dividing the cells into two cluster with the following parameters init_clust=”None”, frac_init_clust=0.5, iter_init=10, doublet_prior=0.5.24 In addition, for the purpose of validation the discriminating SNP list generated by freemuxlet was clustered with SNPs called by samtools using the same common SNP database from two samples of entirely maternal (blood) or fetal (cord blood) origin (**Fig. S10A-S10C**).

### Chromosomes X and Y FISH library design and synthesis

We selected one 1-Mb region for each of human chromosomes X and Y (T2T chrX: 10000000-11000000; T2T chrY: 2781480-3781479), which were previously known to exclude pseudoautosomal regions. (51) (52) These sequences were split into 1-kb fragments, and were then processed using OligoArray2.1 (53) to identify candidate FISH oligonucleotide sequences, using the following parameters: 30 nt oligo length, without overlap; melting temperature between 65°C and 85°C; GC content between 30% and 90%; no secondary structure with a melting temperature greater than 70°C; no cross-hybridization among oligos with a melting temperature greater than 70°C; and no single-nucleotide repeats of length 6 or more. OligoArray2.1 was additionally provided with a FASTA file containing a single sequence of ∼70 kb “A” nucleotides for its built-in specificity filtering step. Candidate oligos were filtered for binding specificity in two steps: (1) filtering out oligos with a match score of 33 or more using BLAST+ blastn-short (54) against the repetitive sequences of the human genome (Repbase (55) [https://www.girinst.org/repbase/, Repeat class: “All”, Taxon: “Homo sapiens (Human)”, including ancestral]); (2) filtering out any remaining candidate oligos that have more or less than one BLAST+ megablast (56) match against the unmasked human genome (T2T). From the remaining candidate sequences, we selected 500 for each chromosome that resulted in the highest probe densities along the linear genome. A 20-nt readout sequence was concatenated to each end of the selected FISH probe sequences (Table 2), with a single “A” nucleotide spacer between all sequence components. Finally, a 20-nt primer binding sequence was added to each end of all sequences for PCR-based library amplification. FISH probe libraries were synthesized from template libraries using to a previously published protocol. (57)

**Table 2.**
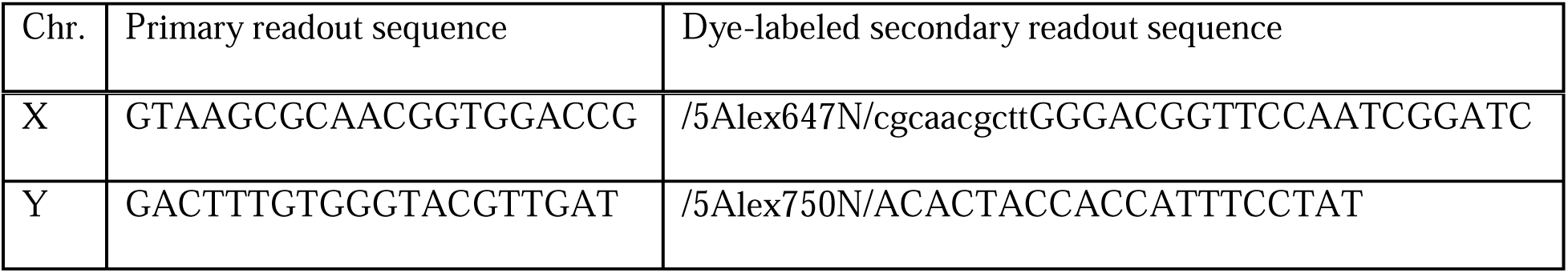
Primary and secondary readout sequences for chromosomes X and Y FISH. Lowercase nucleotides are not utilized in adapter construction.

### Sample preparation for imaging

Glass coverslips (40-mm-diameter, #1.5, Bioptechs, 40-1313-03193) were washed in 50:50 37% hydrochloric acid and methanol at room temperature (RT) for 30 min, rinsed in Milli-Q water 3 times, rinsed in 70% EtOH once, and dried in a 70°C oven. Coverslips were then silanized for 30 min at RT in chloroform mixed with 0.4% v/v allyltrichlorosilane (MilliporeSigma, 107778) and 0.1% v/v triethylamine (MilliporeSigma, TX1200). Silanized coverslips were rinsed once with chloroform, once with ethanol, and were then dried in a 70°C oven for 1 hr. Next, coverslips were treated with poly-D-lysine (1 mg/mL; EMD Millipore, A-003-E) at RT for 30 min, followed by air drying at RT for 30 min.

Tissue was sectioned in a cryostat (Leica, CM3050 S) at -20°C and 20 µm thickness onto treated coverslips. Samples were allowed to dry at RT for 10 min, and were then fixed for 10 min at RT with 4% paraformaldehyde (PFA; Electron Microscopy Sciences, 15710) in Dulbecco’s phosphate-buffered saline (DPBS; Sigma-Aldrich, D8537), and were then washed with DPBS for 3 times, each for 5 min. Samples were permeabilized for 15 min at RT with gentle shaking in a solution of 1.0% v/v Triton X-100 (Sigma-Aldrich, X100) in DPBS, followed by washing 3 times, each for 5 min, in DPBS. Samples were then treated for 5 min at RT with 0.1 M hydrochloric acid, and were washed 3 times with DPBS, 5min each. Samples were digested with RNase (Invitrogen, AM2286; 1:100) in DPBS for 45 min at 37°C, and were then rinsed 3 times with DPBS.

Samples were treated using an Endogenous Biotin-Blocking Kit (Invitrogen, E21390, and were then incubated for 30 min at RT in blocking buffer consisting of 10% w/v Bovine Serum Albumin (Sigma-Aldrich, A9647), 300 mM glycine (AmericanBio, AB00730), and 0.1% v/v Tween-20 (Sigma, P7949) in DPBS. Primary antibody staining was performed by incubating the sample in anti-HLA-DR antibody (Abcam, ab55152, 1:100) in blocking buffer overnight (>16 hr) at 4°C. After primary antibody staining, samples were washed 3 times, 5 min each, in DPBS, and were then incubated in biotin-conjugated secondary antibody (Jackson ImmunoResearch, 715-065-151, 1:1000) for 1 hr at RT in blocking buffer, followed by 3 washes, 5 min each, in DPBS. After staining, samples were postfixed in 4% PFA in DPBS for 5 min at RT, washed with DPBS 3 times, each for 2 min, and then further fixed with 1.5 mM BS(PEG)9 (Thermo Scientific, 21582) in DPBS for 20 min at RT. Samples were then washed with DPBS 2 times, each for 2 min, and were then quenched with 100 mM Tris PH 7.4 (AmericanBio, AB14044) for 10 min at RT. Finally, samples were embedded in 4% Acrylamide/Bis 19:1 gel (Bio-Rad, 1610144) for 1.5 hr at RT.

For FISH probe hybridization, samples were incubated at RT for 30 min in 50% formamide (Sigma-Aldrich, F7503) and 0.1% v/v Tween-20 in 2xSSC (Invitrogen, 15557044). Samples were then sandwiched against the bottom of a small cell culture dish in a 100 µl buffer consisting of 50% formamide and 20% Dextran Sulfate (EMD Millipore, S4030) in 2xSSC, and containing ∼2,400 ng each of synthesized FISH probe libraries targeting human chromosomes X and Y. Samples were then denatured in a water bath at 83°C for 10 min, and were then incubated at 37°C in a humid chamber for at least at least 36 hr (2 overnights). After FISH hybridization, samples were washed with 0.1% v/v Tween-20 in 2xSSC 2 times, each for 15 min, in a 60°C water bath, and then 1 additional 15 min wash at RT.

For visualization of MHCII staining, samples were each incubated in 3 mL blocking buffer containing 3 µg streptavidin (Jackson ImmunoResearch, 016-540-084), and were then washed in DPBS 3 times, each for 5 min. For visualization of chromosome X and Y FISH, readouts were hybridized to samples by incubation for 30 min at RT in a buffer consisting of 35% formamide and 0.01% Triton X-100 in 2xSSC, containing adapter oligonucleotides (30 nM; containing a docking sequence to the readout sequence of the primary FISH probes, and 2x docking sites for dye-labeled readout oligo binding) and dye-labeled secondary readout oligonucleotides (60 nM) for each of the chromosome targets. Readout sequences are provided in Table 2, above. Dye-labeled secondary readout oligonucleotides and adapters were ordered from Integrated DNA Technologies (IDT). After readout hybridization, samples were washed in 30% formamide in 2xSSC for 5 min at RT with gentle rocking. Samples were incubated with 1:1000 DAPI (Thermo Scientific, 62248) in DPBS, and where then transferred to an imaging buffer consisting of 50 mM Tris-HCl pH 8.0, 10% wt/v glucose, 2 mM Trolox (Sigma-Aldrich, 238813), 0.5 mg/mL glucose oxidase (Sigma-Aldrich, G2133), and 40 μg/mL catalase (Sigma-Aldrich, C30) in 2xSSC (58). Samples were imaged using a home-built microscope system which has been previously described (57). Images were collected at 200 nm increments to a total depth of 18 µm. Chromosomes X and Y FISH, MHCII antibody staining, and DAPI were imaged sequentially at every z-height, within the same image stack, to prevent z-drift among different imaging channels.

### Derivation, culture, and storage of CTBs from human placental villi

Term placental villi tissue was digested with 0.25% trypsin (Gibco 15090046) and 0.5 mg/mL DNase I in Advanced DMEM/F12 medium (Gibco 12634010). CTBs were isolated from digests by collecting the 35-45% percoll fraction following a percoll centrifugation. Contaminating immune cells were removed from the retrieved CTBs using CD45 microbeads selection. Purified CTBs were grown on 6-well plates coated with 5 μg/mL human collagen IV (Humabiologics PCIVL) in 2 mL trophoblast organoid medium (TOM, as described in (35)). Cultures were maintained in 5% CO2 in a humidified incubator at 37 °C. Medium was replaced every 2–3 days. For cryopreservation of CTBs, dissociate cells from plates by adding 1mL TrypLE express (Gibco 12604013) and let incubate at 37°C for 15min. Dissociated CTBs were resuspend in freezing media (10% DMSO in FBS) and let slow freeze in a Mr. Frosty freezing container. Frozen cells were moved to liquid nitrogen for long term storage after 24 h.

### Trophoblast organoids coculture with T cells

Cryopreserved placental villi tissue was transferred into a gentleMACS C Tube (Miltenyi Biotec) with human umbilical cord dissociation kit (Miltenyi Biotec 130-105-737) and ran on the gentleMACS Octo Dissociator with Heaters (Miltenyi Biotec) using program 37C_m_LPDK_1. Dissociated tissue was serially filtered through a 100-μm and a 70-μm cell strainer (Thermo Fisher Scientific). Dead cells were removed from digests using a dead cell removal kit (Miltenyi Biotec 130-090-101) and T cells were isolated using human CD3 microbeads (Miltenyi Biotec 130-097-043). Red blood cells were removed from cord blood using RBC lysis buffer, and the resulting cells were treated with dead cell removal and CD3 microbeads to retrieve cord blood T cells. Purified T cells were mixed with CTBs at 1:10 ratio and resuspended in hESC-Qualified Matrigel (Corning 354277) on ice and plated in one 40 µl drop per well into a 24-well culture plate (CytoOne CC7682-7524). 500 µl trophoblast organoid medium (TOM) was added to each well after polymerization of Matrigel at 37 °C for 15 min. Cultures were maintained in 5% CO2 in a humidified incubator at 37 °C. Medium was replaced every 2–3 days. Brightfield images were obtained using an ECHO Revolve microscope. Organoid sizes were quantified using Fiji.

### Trophoblast organoid immunostaining

TOM was aspirated off the wells containing the TOs. 500 μL of ice-cold cell recovery solution (Corning 354253) was added per well, rocking at 4°C for 10min, to release the organoids. Released TOs were fixed in 4 % PFA in PBS for 45 min at RT. The fixed TOs were washed with immunofluorescence (IF) buffer (0.1 % (w/v) BSA, 0.2 % (v/v) Triton X-100, 0.1 % Tween20 (v/v) in PBS). Antigen retrieval solution (Antigen Unmasking Solution (Vector Laboratories H-3300), 0.1 % Tween20 (v/v) in dH20) was added to fixed TOs and heated at 98°C for 20 min. After then, TOs were resuspended in Perm and Block buffer ((5 % donkey serum to 0.5 % (v/v) Triton X-100 in PBS) and transferred to a 96-well plate (Thermo Scientific 163320) to incubate for 1 h at RT. After blocking, TOs were washed three times in IF buffer. Primary antibodies cocktail diluted in IF buffer (1:1000 Anti-Cytokeratin 19 (R&D Systems AF3506), 1:200 Anti-hCG beta (Abcam EPR29783-529), 1:200 Anti-Ki67 (Abcam ab279653)) were then added to TOs and incubated overnight at 4°C. The next day, TOs were washed three times in IF buffer. Secondary antibodies (anti-sheep-647 (Invitrogen A21448), anti-mouse-555+ (Invitrogen A32773), anti-rabbit-488(Invitrogen A21206)) diluted 1:200 in secondary buffer (10% donkey serum in IF buffer) were added to TOs and incubated for 1 h at RT in the dark. The TOs were washed three times in IF buffer and stained with 1μg/mL DAPI (Invitrogen D1306) for 10 min at RT in the dark. TOs were washed 3 times with PBS and mounted on slides (Fisher brand 12550143) in VectaMount mounting media (Vector Laboratories H-5700-60). The TOs were imaged on an ECHO Revolve microscope.

### Bulk-RNA sequencing of trophoblast organoid

On the last day of co-culture experiment, TOM was aspirated off the wells containing the TOs. 500 μL of ice-cold cell recovery solution (Corning 354253) was added per well, rocking at 4°C for 10min, to release the organoids. RNA of the TOs were extracted using RNeasy Plus Micro Kit (Qiagen 74034) following the instructions of the manufacturer. Purified RNA was sent to Yale Center for Genome analysis for library prep and sequencing services. Returned sequencing results were analyzed using R.

### Statistical Analysis

Pre-processing of single cell transcriptomic data was described above under “Bioinformatics”. Then gene expression data were scaled and normalized in Seurat. False discovery rate of adjusted p-value cutoffs of 0.05 were implemented where possible to correct for multiple testing in large datasets. Statistical tests used in figures are described in the accompanying figure legend.

### Visualization

Plots were generated using Prism v10 (GraphPad), ggplot2 and Seurat in R, or Claude (Anthropic). Experimental schematics were created in BioRender.

## List of Supplementary Materials

Table 1

Figs. S1 to S14

Movies S1 to S2

## Acknowledgments

The authors acknowledge the contributions of the Yale Neonatal-Perinatal Medicine NOuRISH team, Yale University Reproductive Sciences Biobank staff, Keck Microarray Shared Resource staff, Yale Center for Genome Analysis staff, Andrew Martins, Gisela Gabernet, and all members of the Konnikova, Megli, and Yimlamai labs.

## Funding

National Institutes of Health grants R01AI171980 & P01AI179570 (LK)

Yale Program for the Promotion of Interdisciplinary Science (LK)

The Hartwell Foundation (TAR)

National Center for Advancing Translational Science CTSA training grant TL1 TR001864 (TAR)

The contents of this publication are solely the authors’ responsibility and do not necessarily represent the official views of NIH.

## Author contributions

Conceptualization: TAR, CJM, LK

Sample acquisition: CJM, LK

Methodology: TAR, YC, MIR, TAS, SS, DY, JSDR, CJM

Validation: TAR, YC, MIR

Data Analysis: TAR, YC, LB, WG

Writing - Original Draft: TAR

Writing - Review & Editing: TAR, YC, LK

Visualization: TAR, YC

Supervision: SW, RM, JST, CJM, LK

Funding acquisition: CJM, LK

## Data and materials availability

All data and code are available at https://doi.org/10.5281/zenodo.18765352. Inquiries about materials should be directed to the corresponding author.

