## Supplementary material for "Excessive immune responses in preterm placental villi compromise trophoblast health": suplm figures

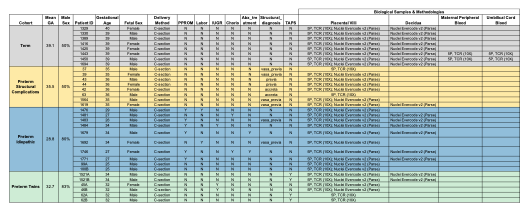

**Table 1. Sample Characteristics and Metadata.** Clinical metadata is provided for each sample (N=32), grouped by study cohort (N=6-10 per cohort): gestational age, fetal sex, delivery method, preterm premature rupture of membranes (PPROM), labor, intrauterine growth restriction (IUGR), chorioamnionitis (Chorio), antibiotic treatment (Abx_treatment), structural diagnosis (vasa previa, placenta previa, placenta accreta), and Twin Anemia Polycythemia Sequence (TAPS). Methodologies used to analyze each sample are also described: 10X Genomics Chromium assays (5P gene expression and VDJ_TCR), and Parse Evercode v2 nuclei assay.

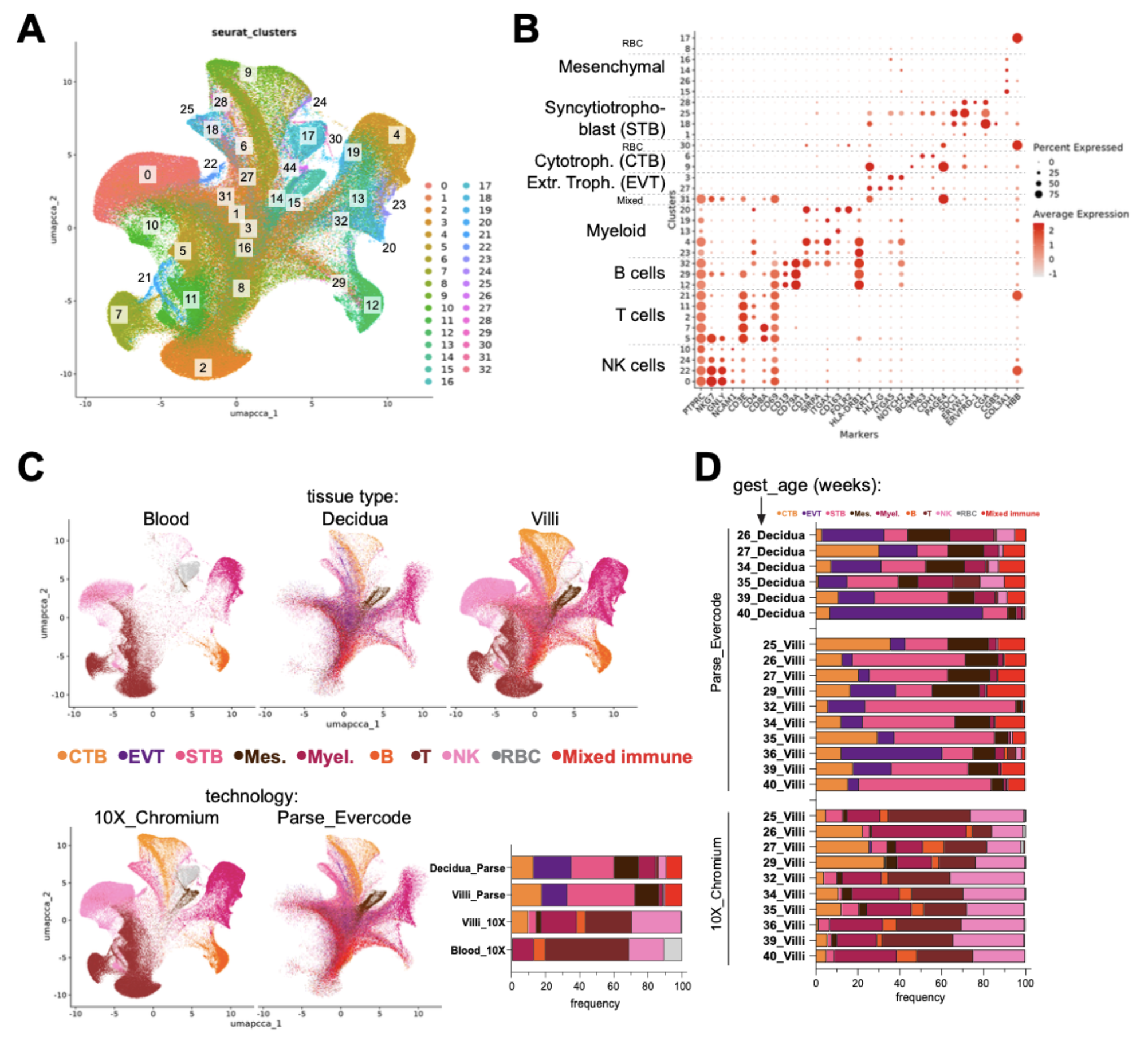

Fig. S1. Annotation of major cell clusters. (A) Unsupervised clustering of integrated placenta immune cell atlas accompanied by (B) marker genes allowing for annotation of cell clusters. (C) UMAP split by tissue type and by technology. Enumerated as stacked bars at bottom. (D) Cell type frequencies across gestational age and technology.

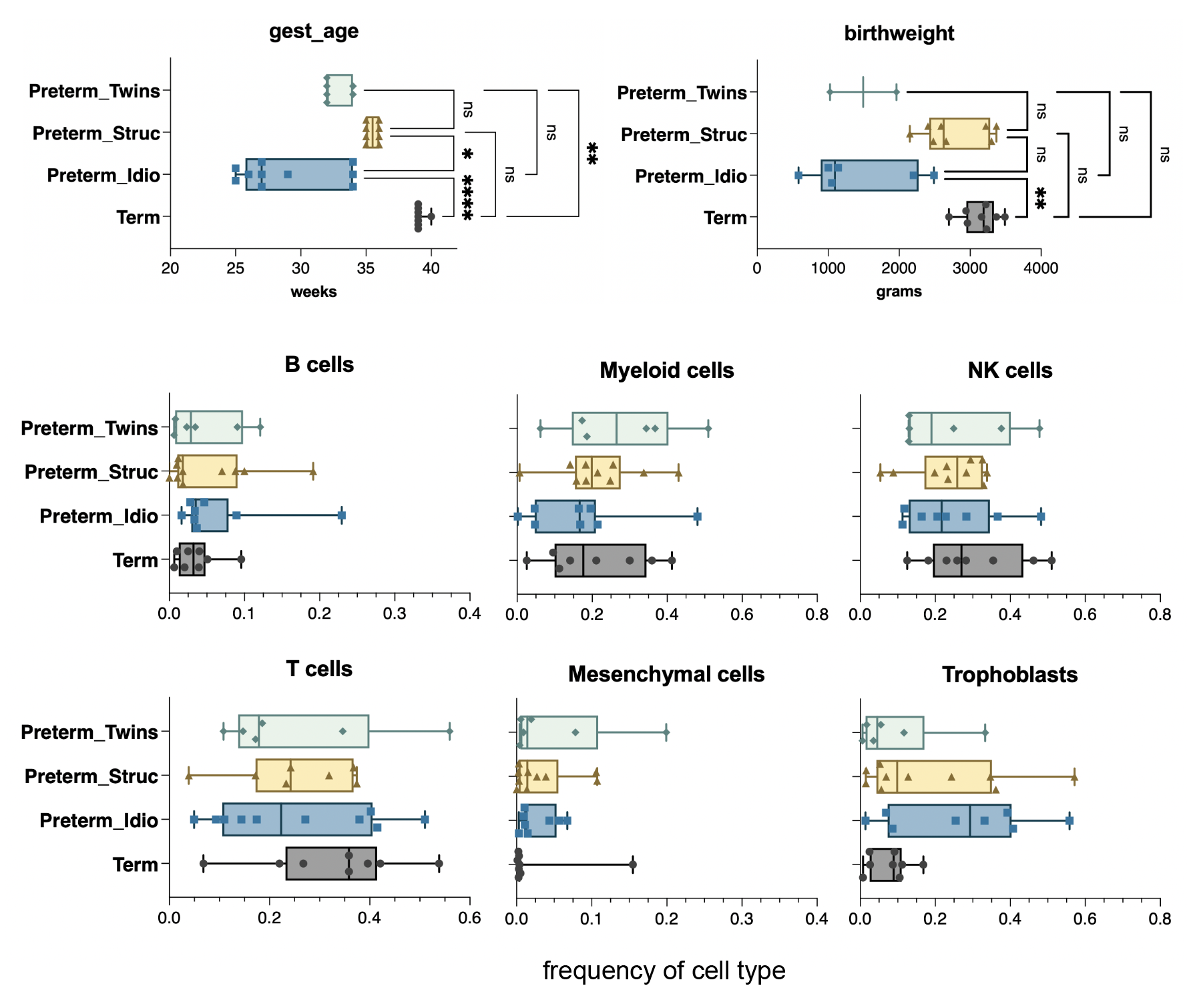
Fig. S2. Major cell types are found across late gestation. (A) Gestational age of placental donors. (B) Birthweight. (C) Frequencies of major cell lineages. N=6-10 per cohort. **P*<0.05, ***P*<0.01, *****P*<0.0001 by Kruskal-Wallis test. ns, not statistically significant.

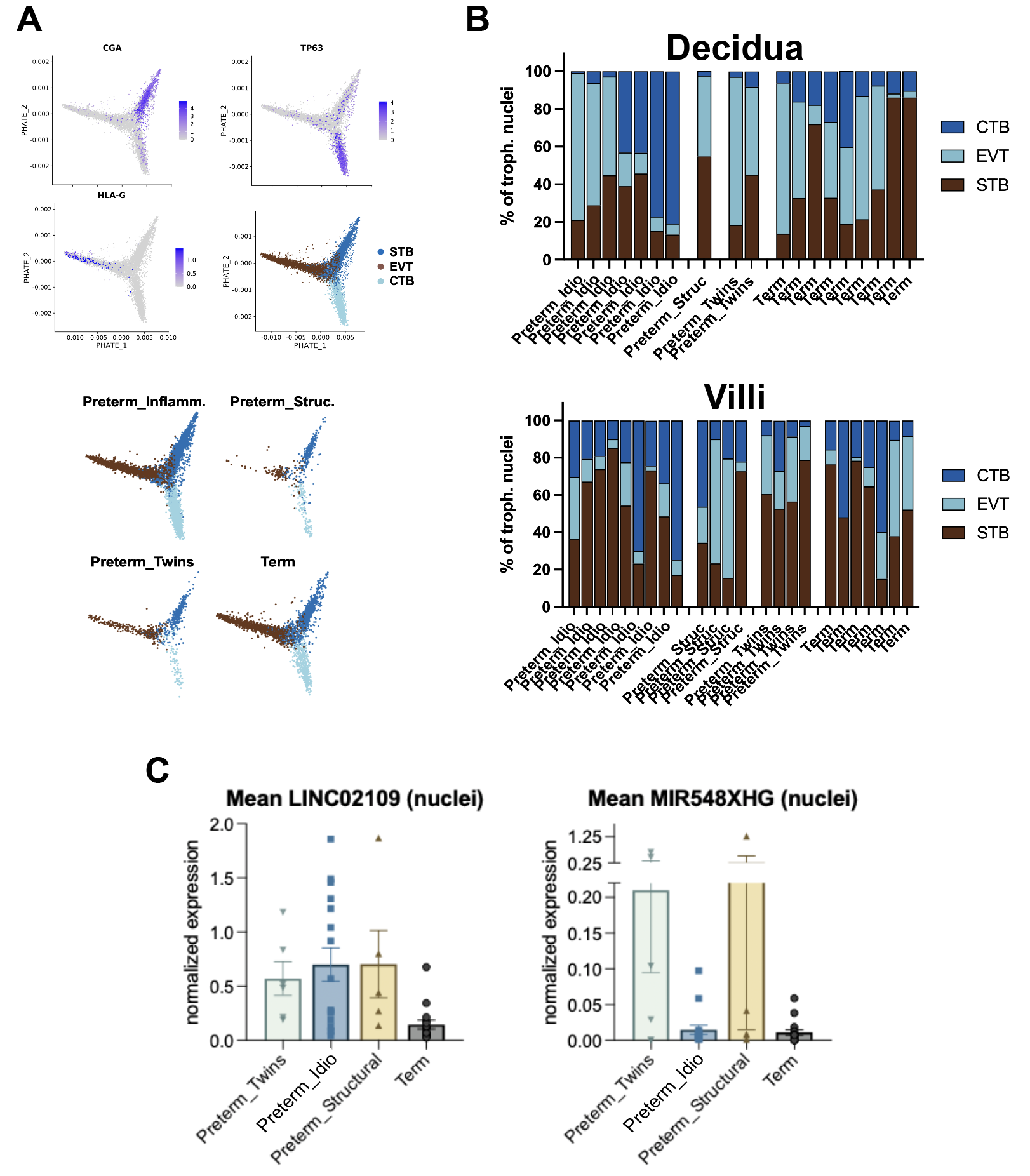

**Preterm_Idio**

**Fig. S3. Trophoblast diversity captured by single nuclei sequencing.** (A) Annotation of major trophoblast clusters based on marker gene expression. Abbrevs: Cytotrophoblasts, CTB; Extravillous trophoblasts, EVT; Syncytiotrophoblasts, STB. (B) Trophoblast population frequencies in individual subjects, separated by tissue type. (C) Differentially expressed genes in trophoblast nuclei from various preterm cases.

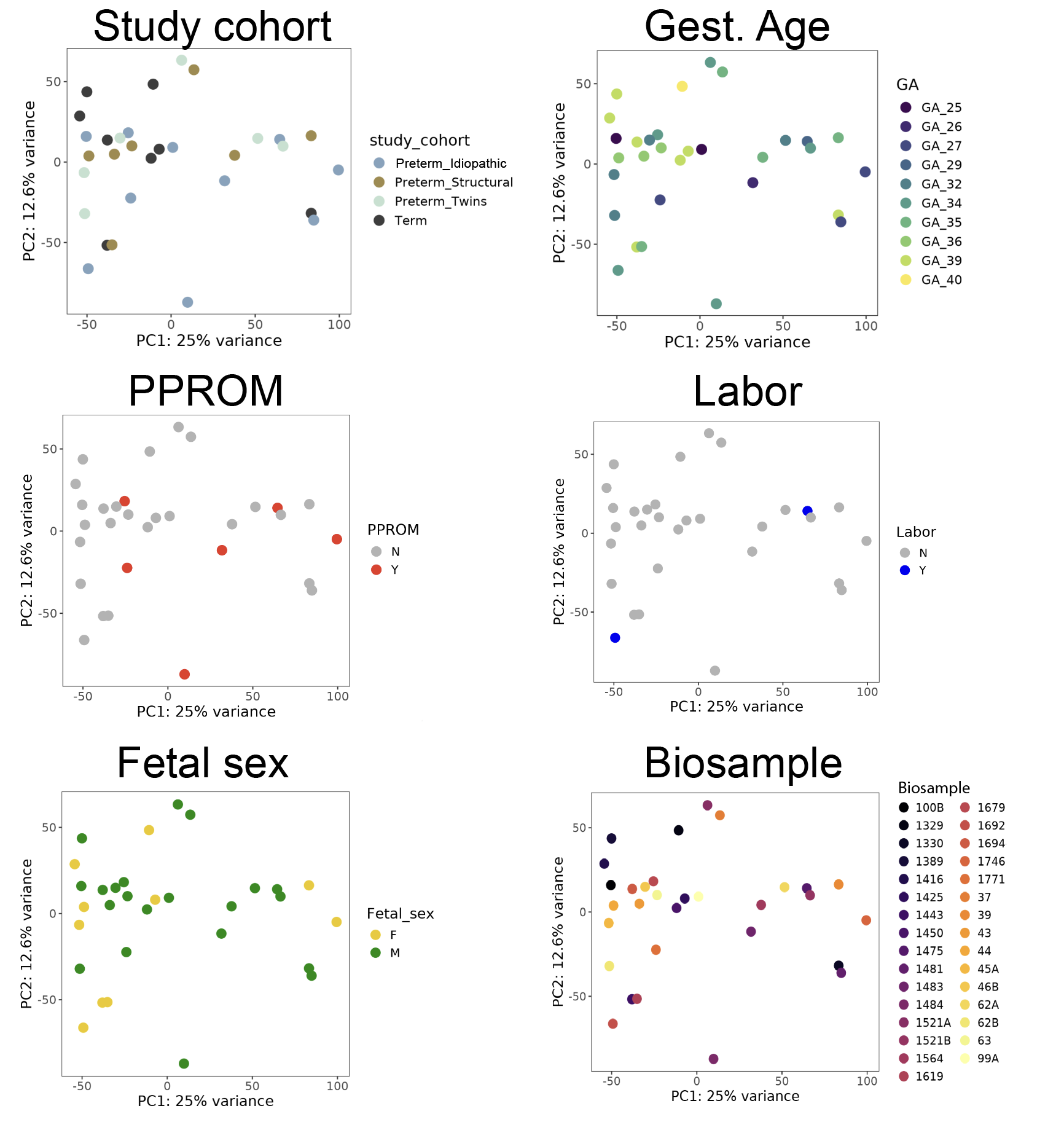

Fig. S4. Pseudobulk analysis across major metadata categories. `Principal component analysis (PCA) of villi scRNAseq samples, colored by study_cohort, gestational age, PPROM, Labor, fetal sex, or biosample.

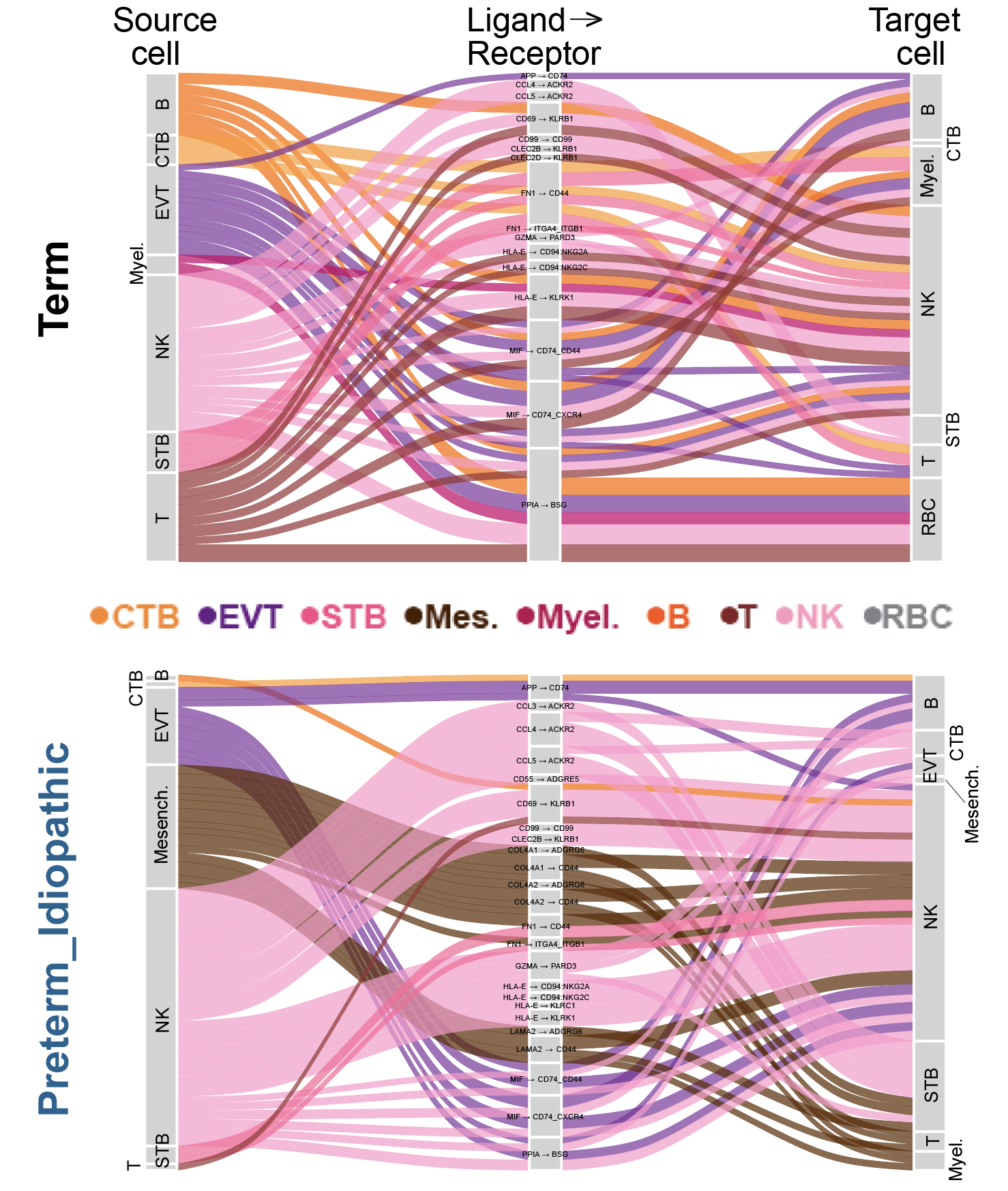
**Fig. S5. Ligand receptor interactions are more diverse in term villi than idiopathic preterm villi.** Cell chat diagrams from term cases and idiopathic preterm cases, with imputed ligand-receptor interactions in the center column between source cells and target cells. Line thickness represents predicted interaction probability.

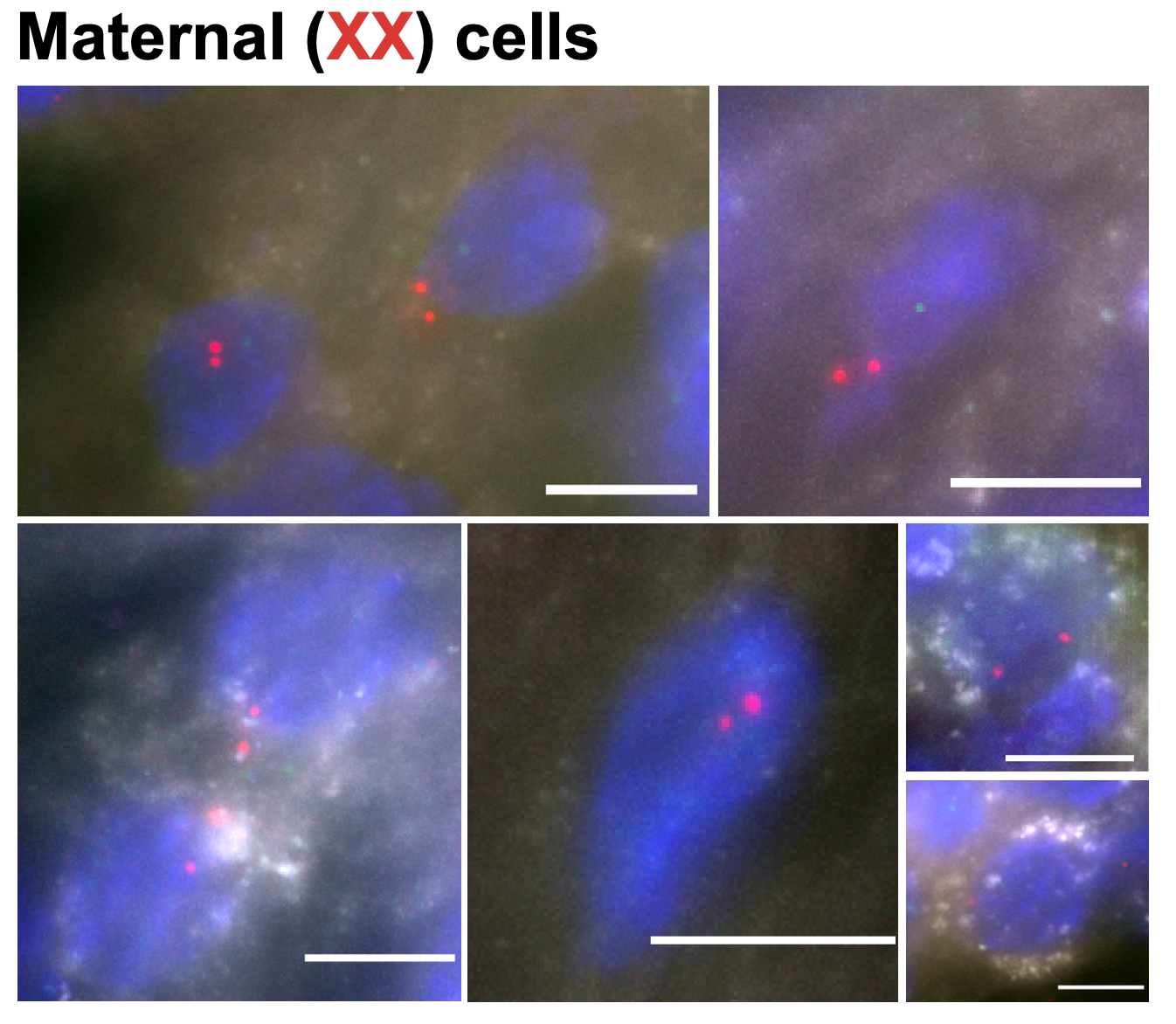

**Figure S6. Heterogeneous HLA-DR expression by maternal (XX) cells in placental villi.** Insets of selected maternal cells expressing two Xchr FISH signals (red) and variable HLA-DR (white). Scale bars, 10µm.

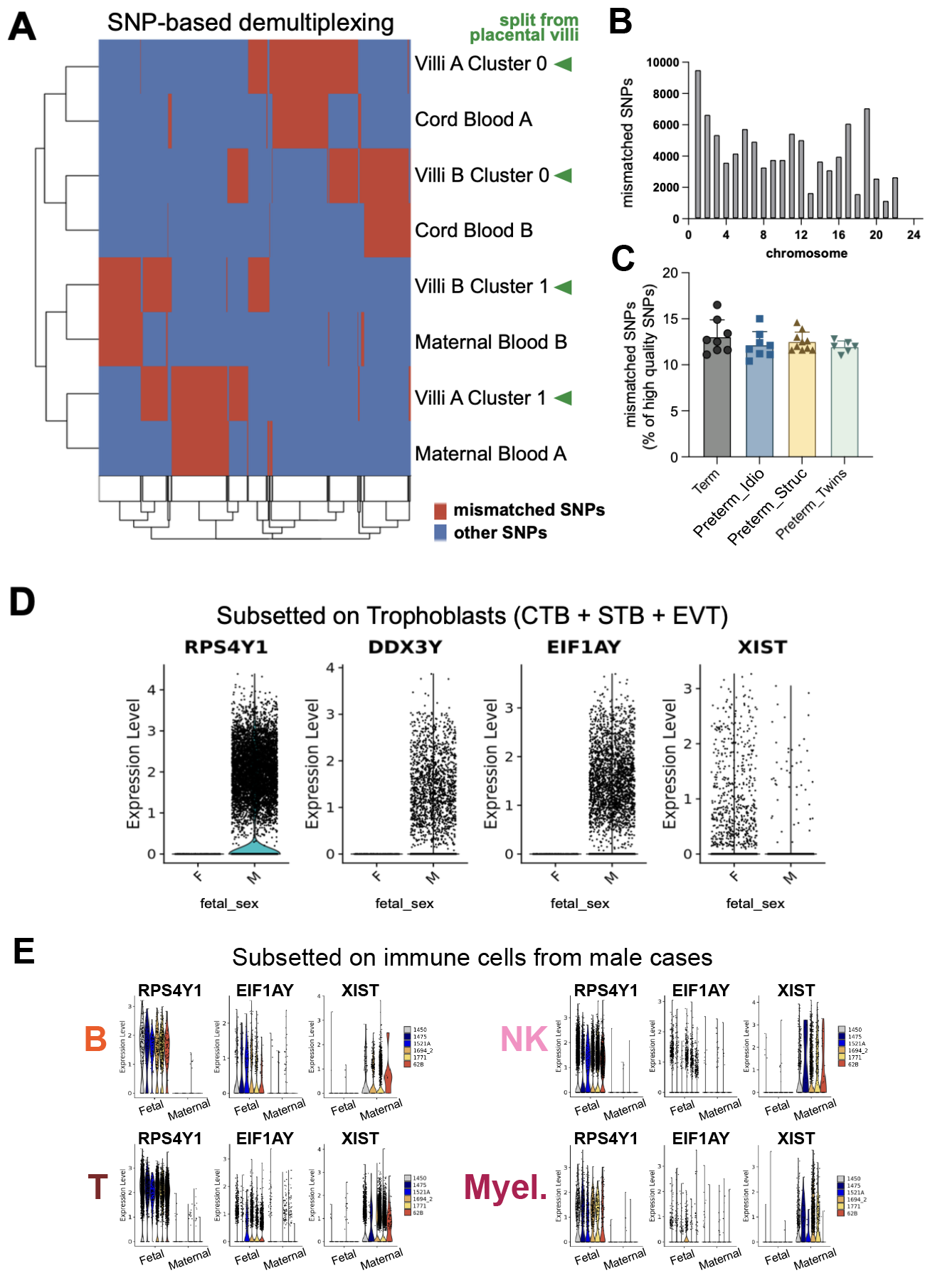
**Figure S7. SNP-based demultiplexing allows for dissection of fetal placental cells from maternal placental cells.** (A) After calling SNPs and filtering on high quality SNPs (≥Q30) that exhibited mismatches, each cell from placental villi was assigned to Cluster 0 or Cluster 1 (green arrowheads), which were then subjected to unsupervised hierarchical clustering with matched maternal blood and cord blood cells and their SNPs. (B) Mapping of mismatched SNPs to their chromosomal locations. (C) Frequency of mismatched SNPs within each dyad across study cohorts. (D) Expression of Y chromosome genes and X chromosome genes in male and female trophoblasts, respectively, or in (E) freemuxlet-separated fetal and maternal immune cells from male cases.

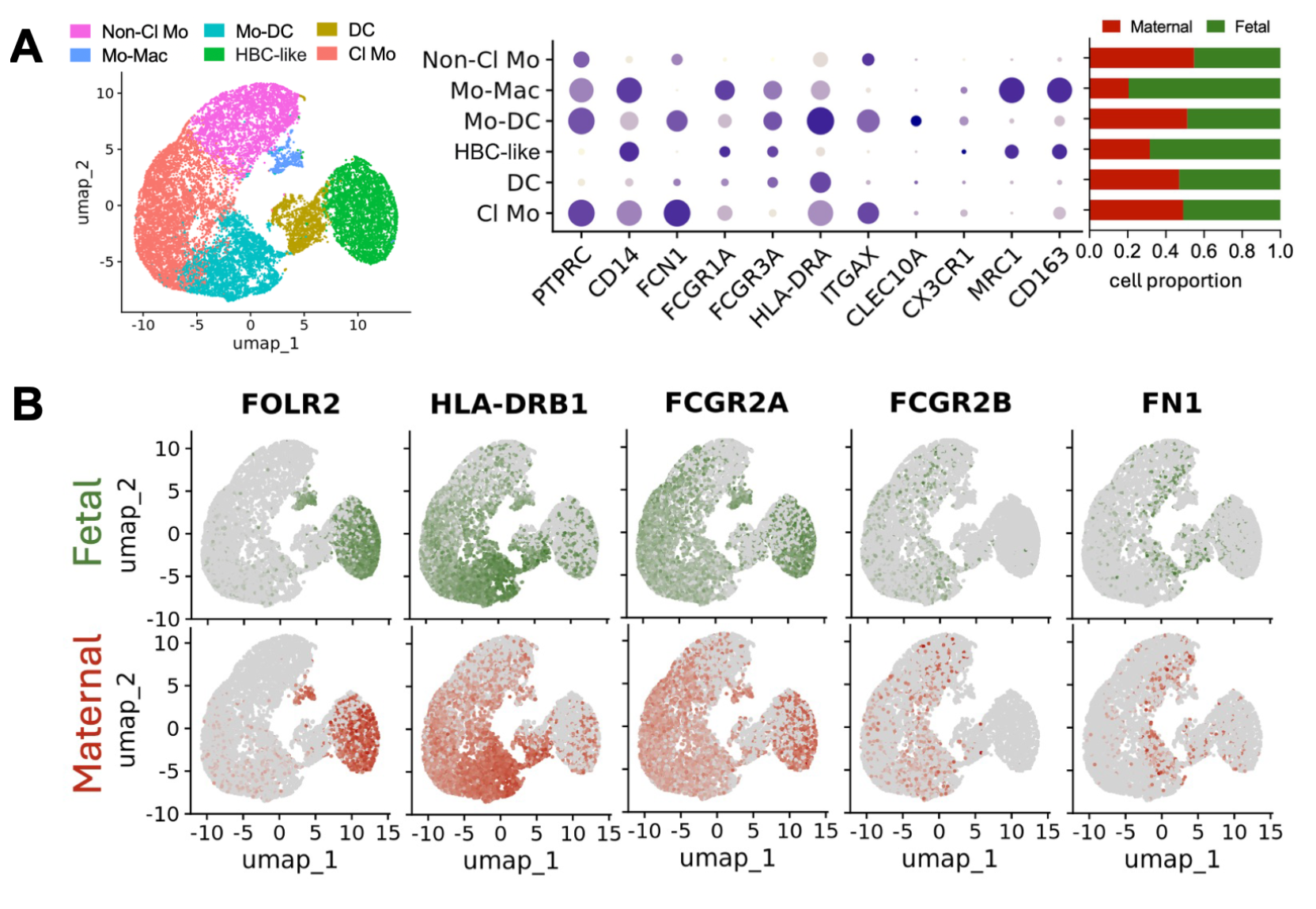

**Figure S8. A spectrum of myeloid cells are associated with structural complications of pregnancy.** (A) UMAP projection of subsetted myeloid cells with annotations, marker gene expression, and frequencies of maternal/fetal origin. (B) Marker genes used to differentiate HBCs and “PAMMs” are broadly present and highly similar across cell origin.

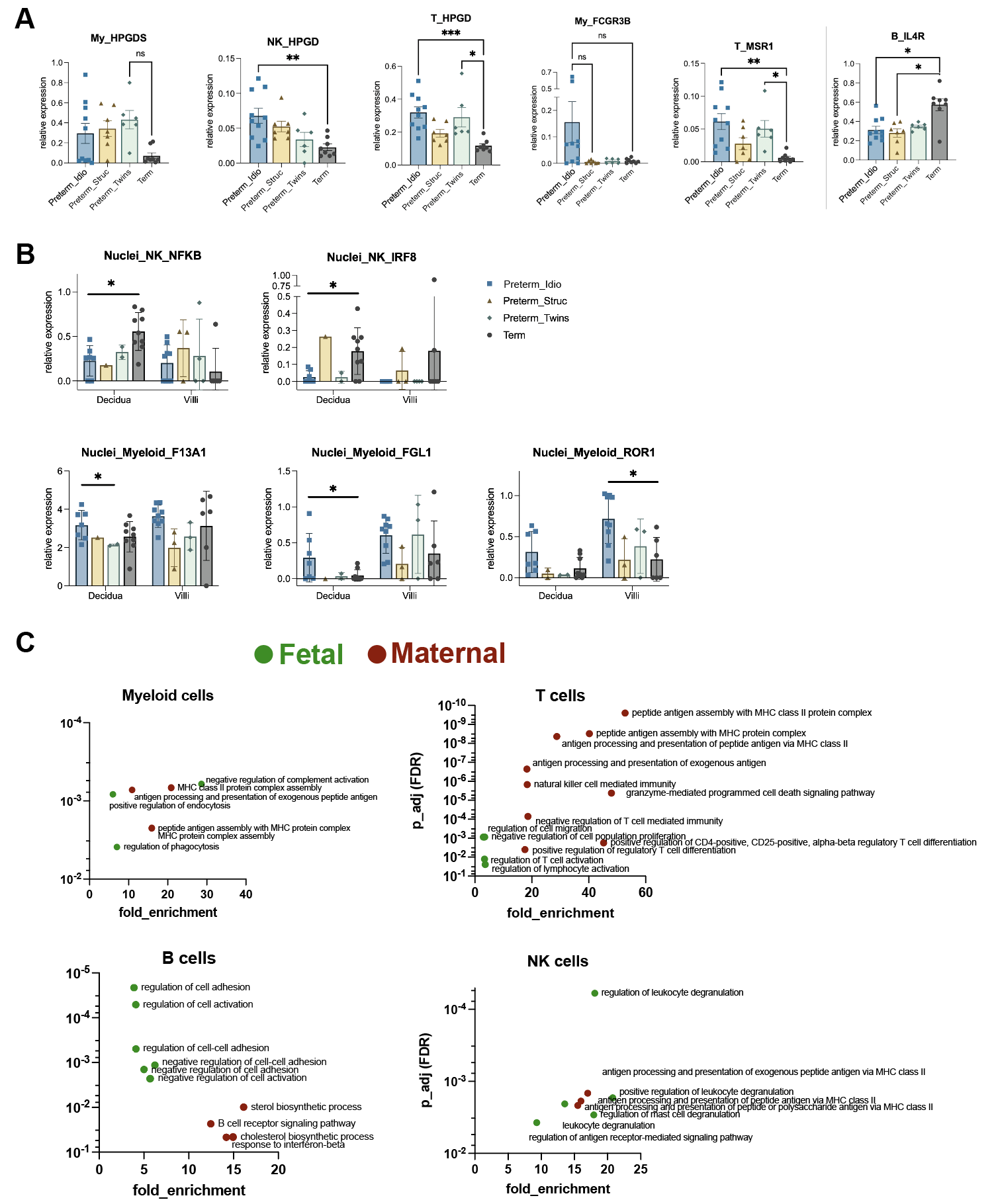

**Figure S9. Prostaglandin D pathways are enriched in fetal immune cells.** (A) Expression of Prostaglandin D Synthase-encoding HPGDS and Hydroxyprostaglandin Dehydrogenase-encoding HPGD in various fetal immune cell subsets. (B) DEGs in decidua and villi nuclei across cohorts. (C) Gene ontology pathways imputed to fetal cells versus maternal cells.

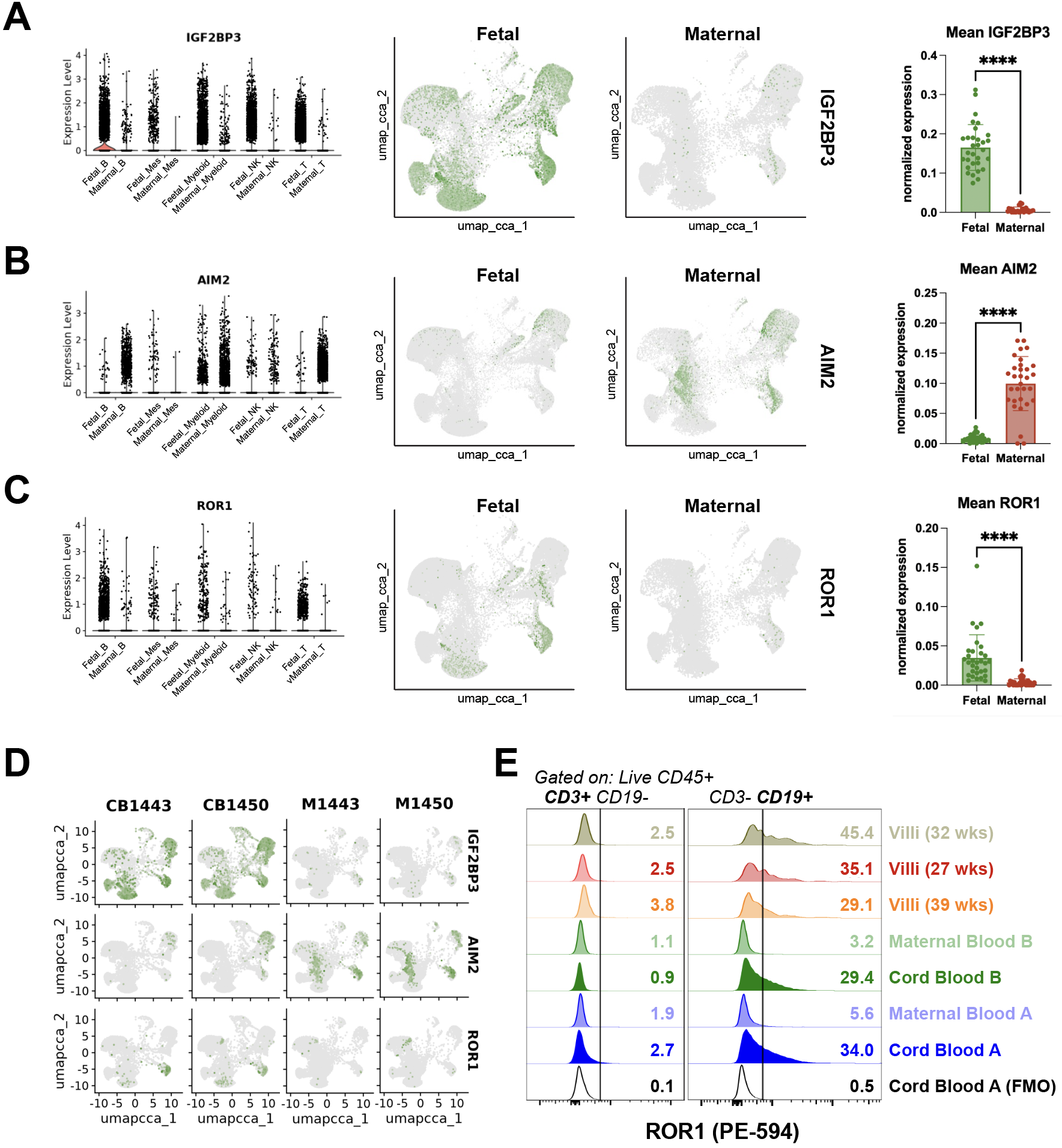

**Figure S10. Identification of candidate fetal-specific and maternal-specific marker genes.** (A) Violin plots split by cell type and UMAP projections of fetal-restricted expression of *IGF2BP3,* maternal-enriched expression of *AIM2* (B), and fetal-enriched expression of *ROR1* (C). ****p<0.0001 by Wilcoxon signed-rank test. (D) Each of the same marker genes of interest plotted in two dyads of cord blood and maternal blood. (E) FACS assessment of surface ROR1 expression in cord blood, maternal blood, and placental villi immune cells.

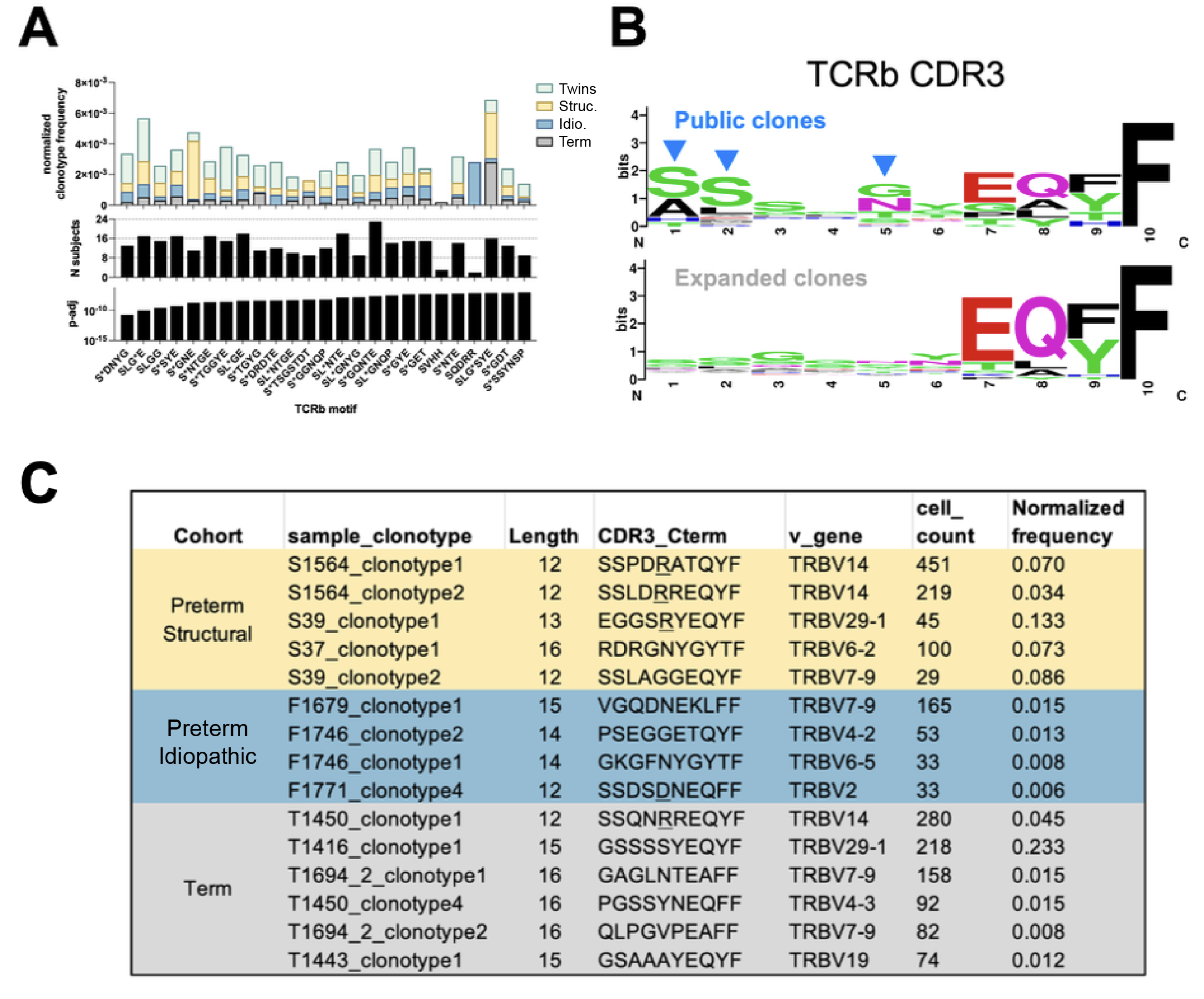

**Figure S11. Public TCR clones have features distinct from the most expanded clones.** (A) Distribution of public clones across study cohorts and their sequence motifs. (B) Sequence logos of  TCR beta CDR3 from top public clones and top expanded clones. Abbrevs: TCRb: T cell receptor beta; CDR3: complementarity determining region 3. (C) Table of most expanded TCR clones, their molecular characteristics, and their frequencies.

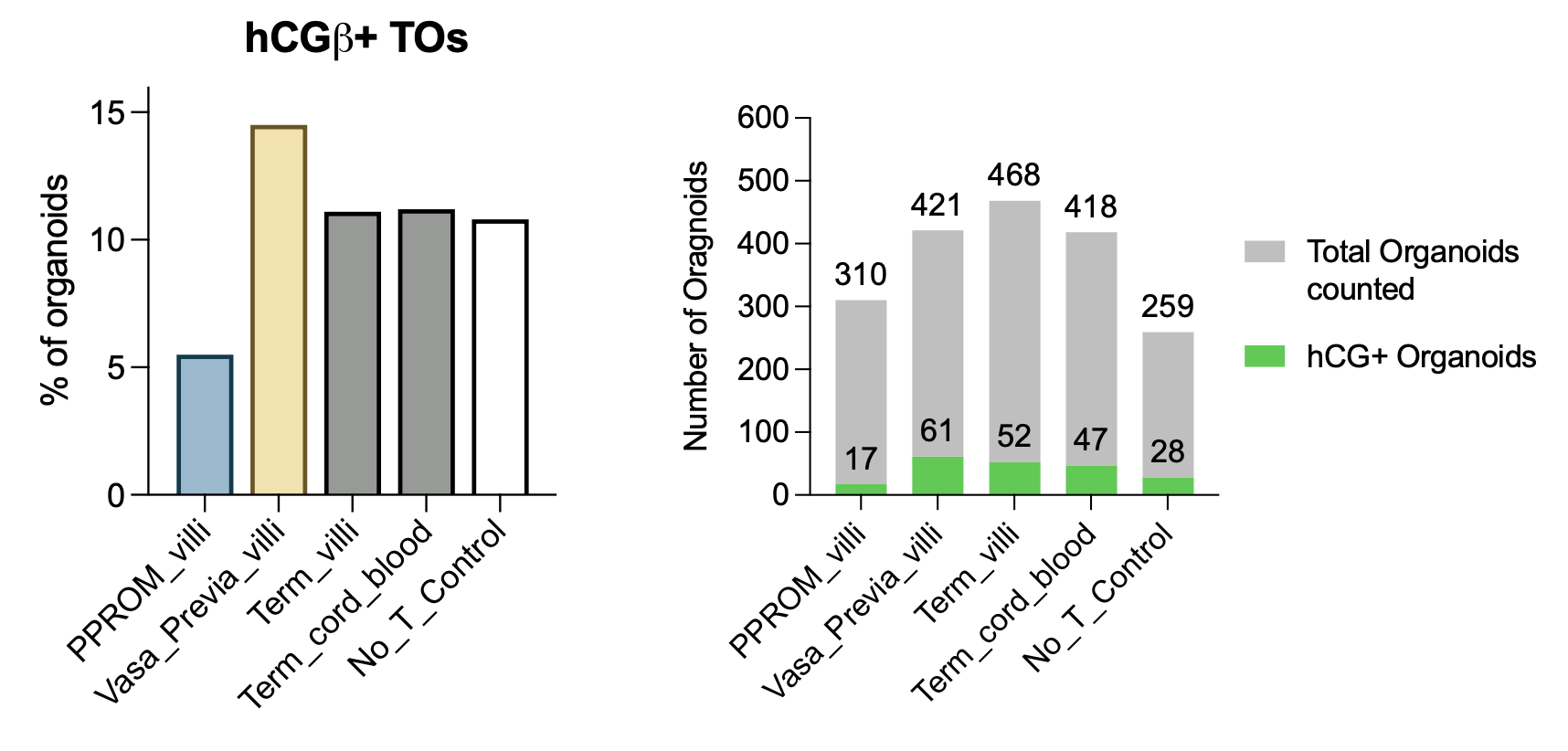

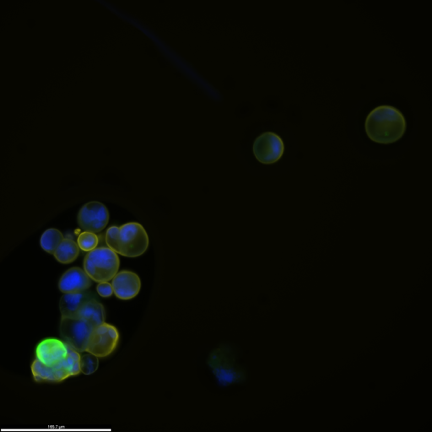

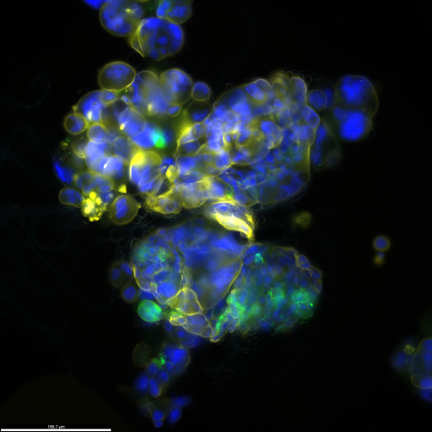

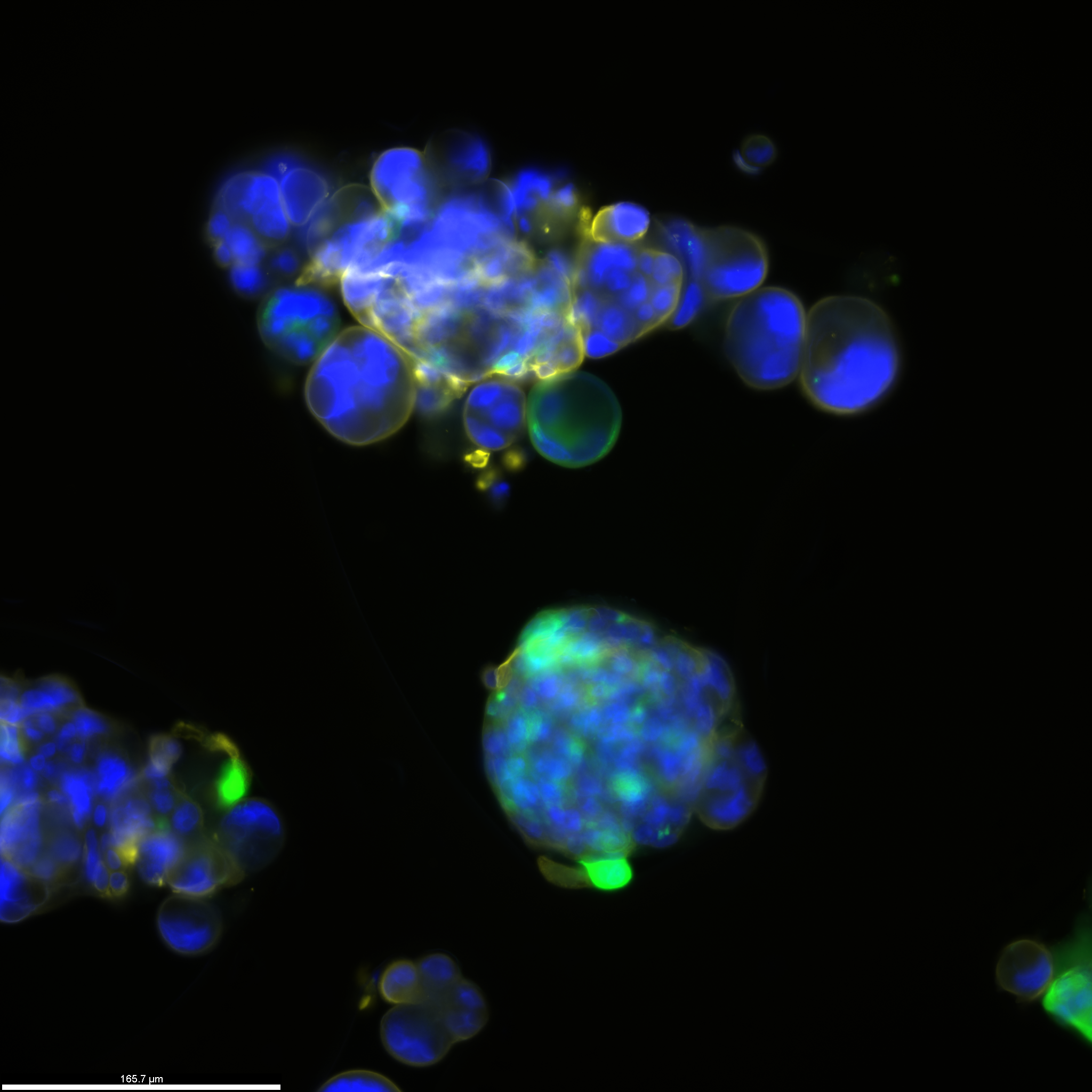

DAPI CK19 hCGβ

PPROM_villi	Term_villi	No_T_Control

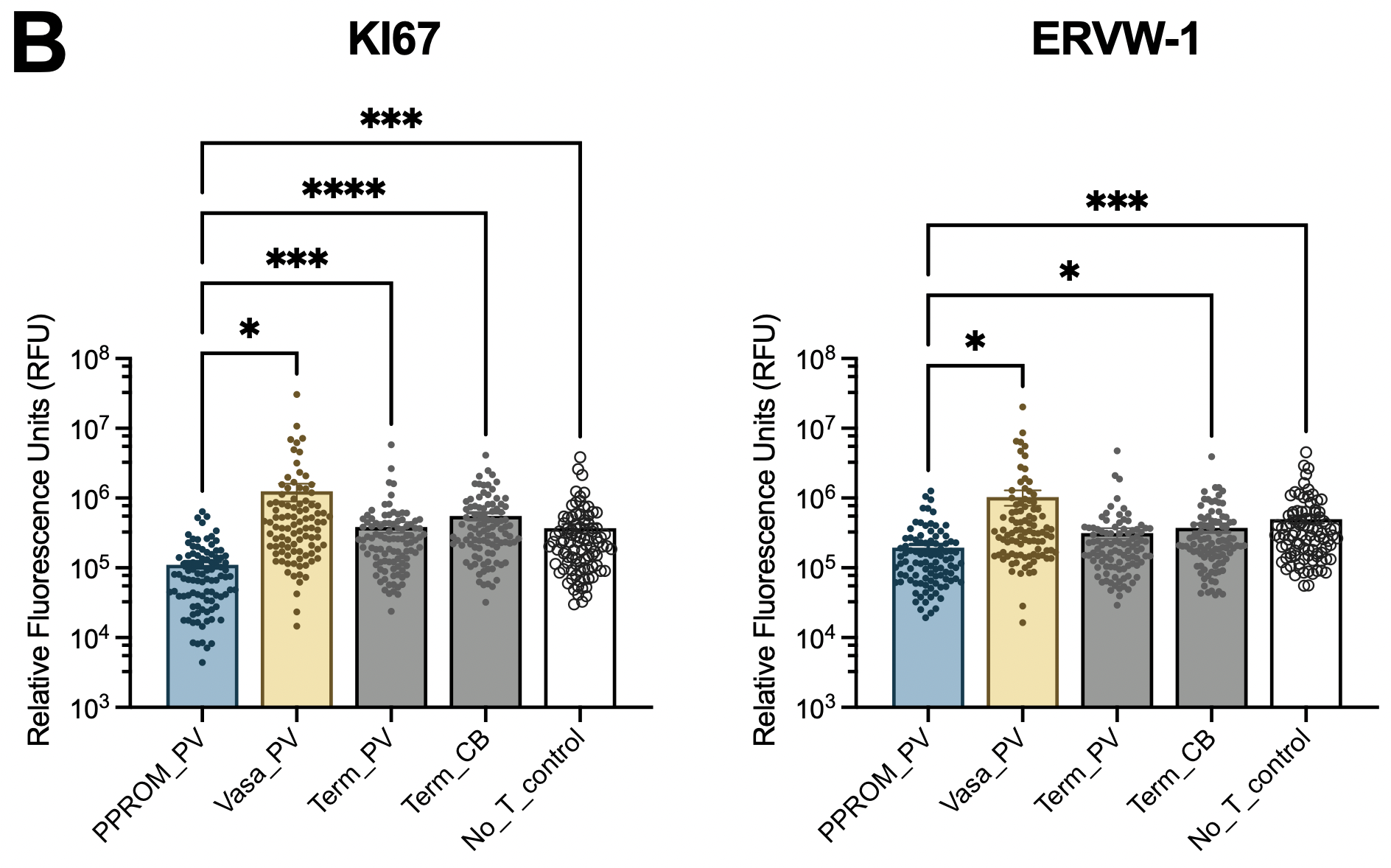

**A**

**Figure S12. Decreased hCG-β expression in trophoblast organoids (TO) co-cultured with PPROM PV T cells.** (A) Immunofluorescence microscopy of hCGβ producing TOs and quantification across co-culture conditions. (B) Quantification of KI67 and ERVW-1 microscopy, such as representative images shown in Figure 8.

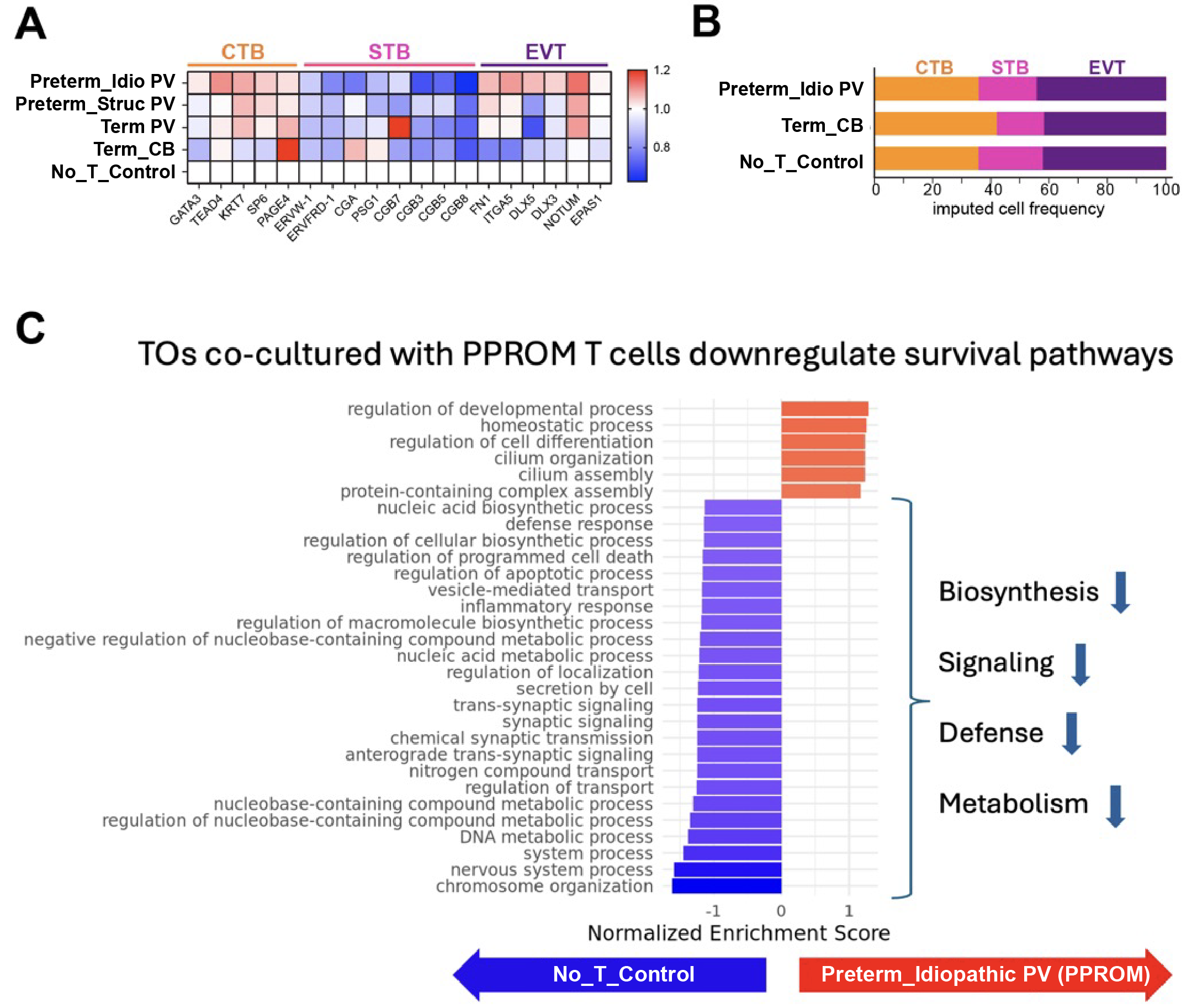

**Figure S13. Trophoblast organoids (TO) contain several trophoblast lineages, and are challenged by co-culture with idiopathic preterm PV T cells.** (A) Trophoblast lineage marker expression across co-culture conditions, using bulk RNAseq. (B) Cell type deconvolution. (C) Gene ontology pathway analysis of idiopathic preterm co-cultures versus No_T_Control organoid cultures.

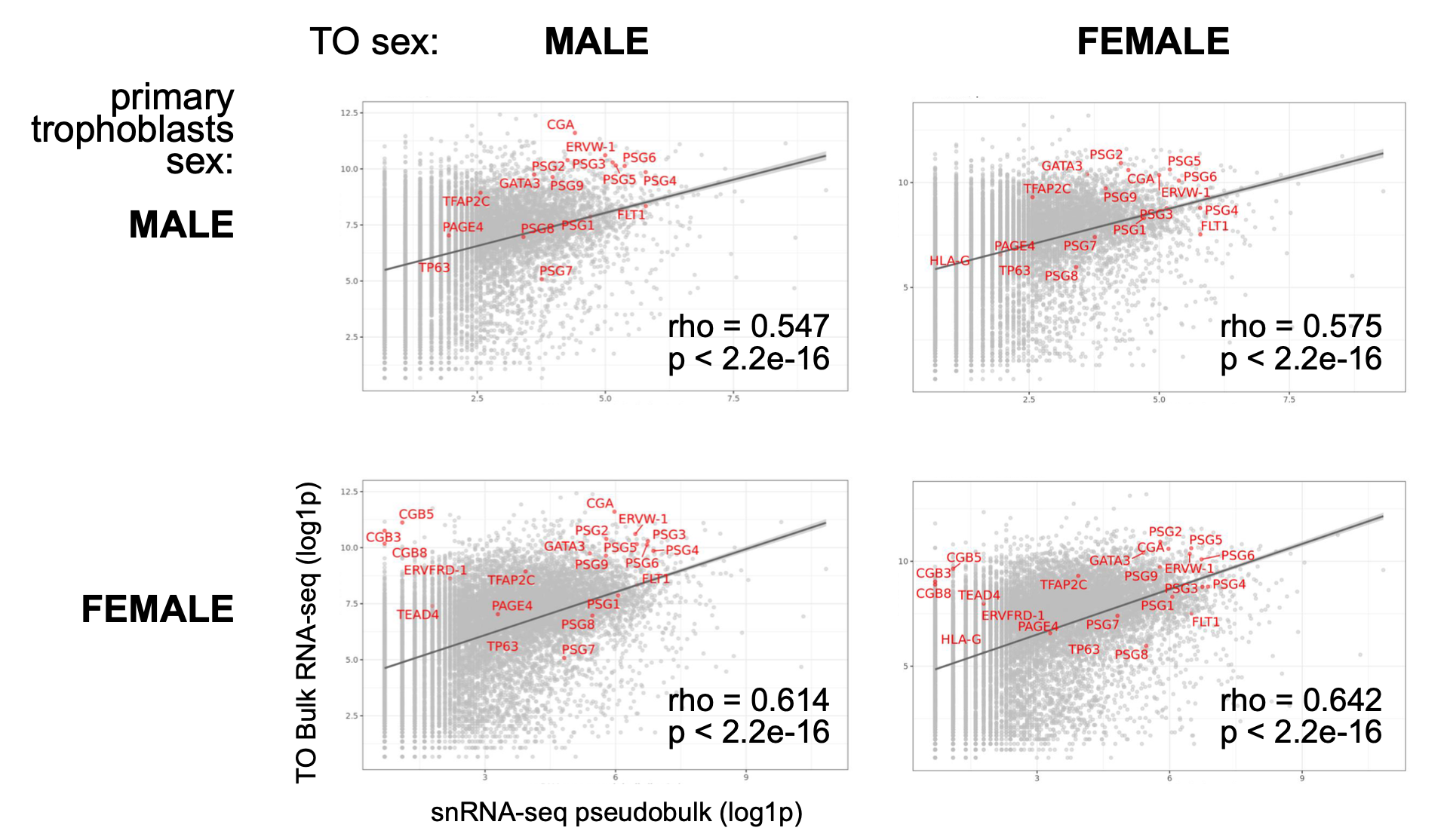

**Figure S14. Correlation between trophoblast organoids (TO) and fresh primary trophoblasts transcripts are not significantly different across fetal sex.** TO bulk RNAseq and fresh primary trophoblast single nuclei RNAseq (snRNAseq) were assessed using Pearson’s product-moment correlation on log(x+1) transformed expression values.

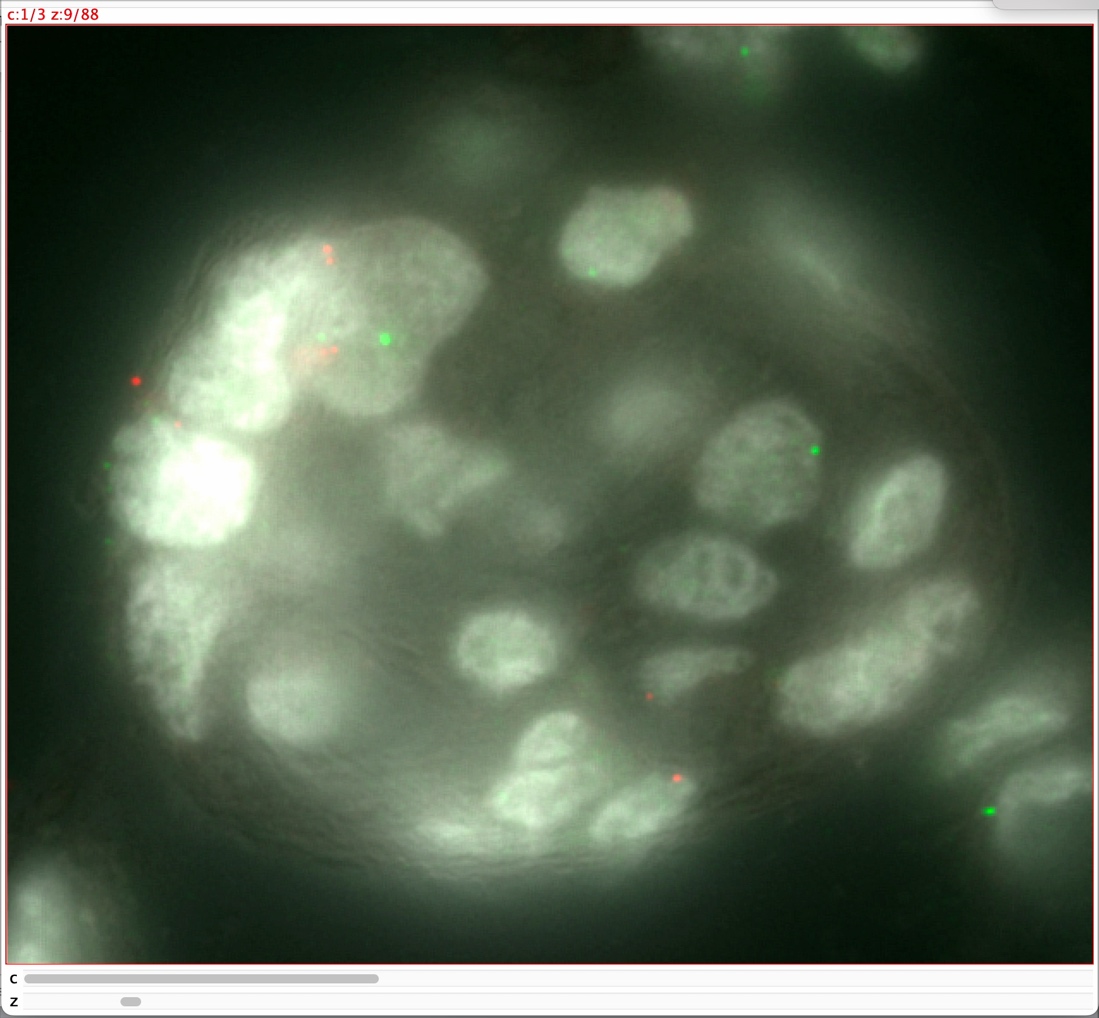

Movie S1. Z-stack flythrough of a villus core of a male placenta, containing XX+ Maternal cells. Ninety (90) images were collected at 200nm z intervals in three channels, DAPI (white), Xchr (red), and Ychr (green), and then stacked in ImageJ. A cell with two consecutive Xchr signals can be seen in the upper middle section of the villus near the end of the movie.

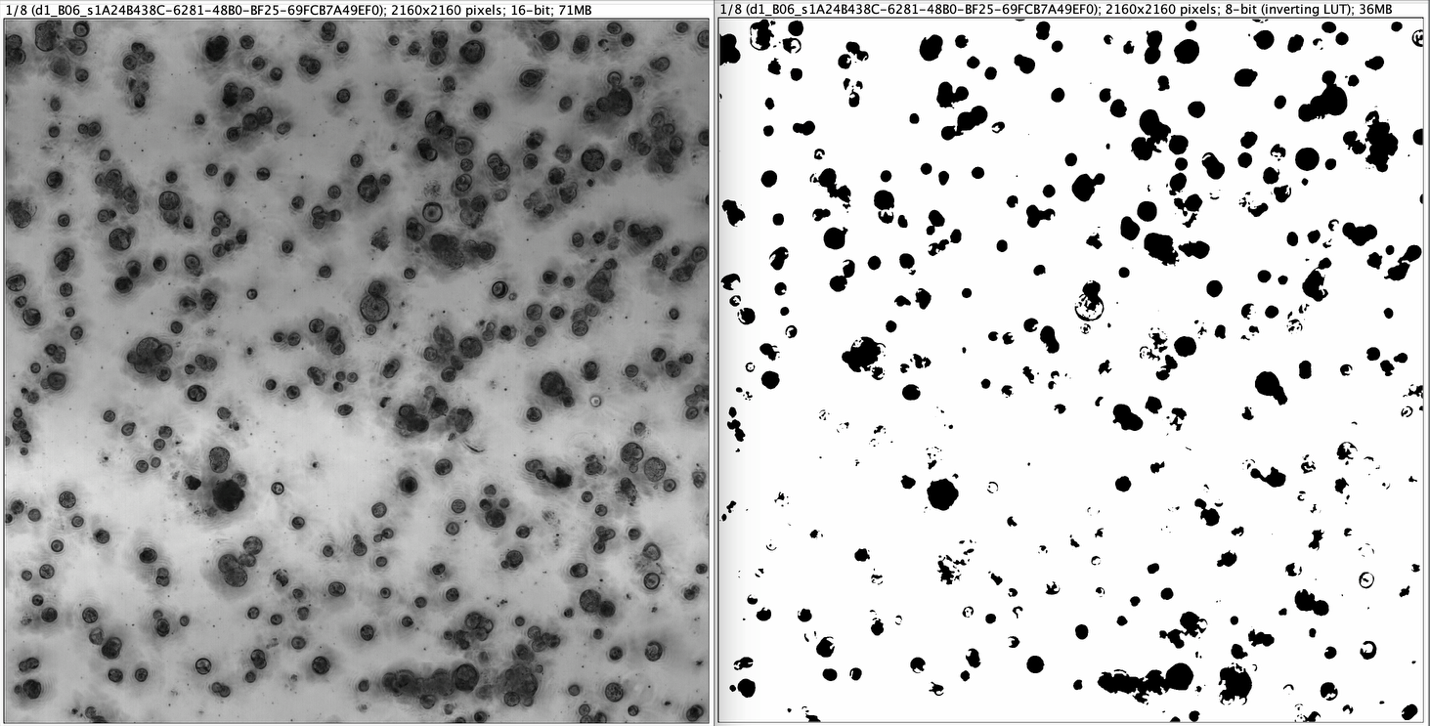

**Movie S2. Time-lapse brightfield micrographs of trophoblast organoid cultures over 8 days.** Brightfield micrographs (left) scale bar, 100µm. Segmentation masks were generated using the TrackMate plugin in ImageJ (right).
